# *Actl7b*-deficiency leads to mislocalization of LC8 type dynein light chains and disruption of murine spermatogenesis

**DOI:** 10.1101/2022.12.19.520998

**Authors:** Gina E. Merges, Lena Arévalo, Keerthika Lohanadan, Dirk G. de Rooij, Melanie Jokwitz, Walter Witke, Hubert Schorle

## Abstract

Actin-related proteins (Arp) are classified according to their similarity to actin and are involved in diverse cellular processes. *ACTL7B* is a testis-specific Arp and highly conserved in rodents and primates. ACTL7B is specifically expressed in round and elongating spermatids during spermiogenesis. Here, we have generated an *Actl7b*-null allele in mice to unravel the role of ACTL7B in sperm formation. Male mice homozygous for the *Actl7b*-null allele (*Actl7b-/-*) were infertile, while heterozygous males (*Actl7b+/-*) were fertile. Severe spermatid defects such as detached acrosomes, disrupted membranes and failed elongation of the axoneme start to appear at spermiogenesis step 9 in *Actl7b-/-* mice, finally resulting in spermatogenic arrest. Abnormal spermatids, were degraded. Co-immunoprecipitation experiments identified interaction between ACTL7B and the LC8 dynein light chains DYNLL1 and DYNLL2, which are first detected in step 9 spermatids and mislocalized when ACTL7B is absent. Our data unequivocally establishes that mutations in ACTL7B are directly related to male infertility, pressing for additional research in men.

**Summary statement:** In this study, Actl7b-deficient mice were generated. Loss of Actl7b leads to spermatogenic arrest in mice. ACTL7B interacts in with DYNLL1/DYNLL2 and seems to function in spermatid cytoskeleton.

## Introduction

Proteins belonging to the superfamily of Actin-like/Actin-related proteins share up to 60% amino acid identity with conventional actins and act in various cellular processes including vesicle trafficking, chromatin modulation, microtubule motility and actin filament dynamics (Schafer and Schroer, 1999).

*ACTL7B* was first described by Chadwick *et al*. (Chadwick et al., 1999), who identified and characterized two novel actin-like genes, *ACTL7A/ T-ACTIN 2* and *ACTL7B/ T-ACTIN 1*, from the familial dysautonomia candidate region on chromosome 9q31 in human. However, neither gene was found to be mutated in patients with dysautonomia suggesting that they are not involved in its pathogenesis. Nucleotide alignment showed high level identity between these two genes and a greater than 40% predicted amino acid identity to a variety of actin proteins. *ACTL7B* is encoded as an intronless gene (Hisano et al., 2003). In head to tail formation *ACTL7A* maps around 6.4 kb apart. In mice *Actl7a* and *Actl7b* were mapped to chromosome 4 (Chadwick et al., 1999, Hisano et al., 2003). It has been proposed these genes arose prior to the divergence of rodents and primates by retroposition of a spliced mRNA transcribed from an actin progenitor gene (Hisano et al., 2003). In mice and human, *ACTL7B* is expressed exclusively in the testis (Tanaka et al., 2003, Hisano et al., 2003), suggesting a role in spermatogenesis.

A study comparing the expression of testis-enriched genes in fertile and teratozoospermic men identified *ACTL7B* to be among those genes significantly lower expressed in teratozoospermic men (Ahn et al., 2017).

In mice and humans, *ACTL7B* has been found to be expressed post-meiotically in round and elongating spermatids (Hisano et al., 2003, Guo et al., 2018). ACTL7B is detected in the cytoplasm and to lesser amounts in the nucleus of round and elongating spermatids and seems to be, in contrast to ACTL7A, evicted with excess cytoplasm at the end of spermiogenesis (Tanaka et al., 2003). Interestingly, 5 polymorphisms in *ACTL7B* (and 6 in *ACTL7A*) were detected in a cohort of Japanese infertile male patients, suggesting that *ACTL7B* plays a role in fertility (Tanaka et al., 2019). Noteworthy, a study comparing two groups of Luzhong mutton sheep with different fecundity identified nine genes, among these *ACTL7B* (and *ACTL7A)*, which were associated with reduced litter size (Tao et al., 2021). Further, proteomic and phosphoproteomic analysis of prepubertal and pubertal testis of swamp buffalo identified ACTL7B to be higher abundant and phosphorylated in the pubertal testis, again suggesting a role in spermatogenesis (Huang et al., 2020). Single nucleotide polymorphisms in the coding sequence of *ACTL7B* in infertile men have been reported, however not been directly correlated to male infertility (Tanaka et al., 2007, Tanaka et al., 2019). Additionally, a recent study, using comparative proteomics on human testicular tissue, identified ACTL7B, both on protein and mRNA level, among the six proteins/transcripts with the highest discriminating power of obstructive and non-obstructive azoospermia subtypes (Davalieva et al., 2022).

Although the molecular function of ACTL7B is not known, its immunolocalization suggests a role in cytoskeletal organization and/or protein trafficking during spermatogenesis. We generated an *Actl7b*-null allele using CRISPR/Cas9-mediated gene editing. While *Aclt7b+/-* males were unaffected, *Actl7b-/-* males were infertile showing spermatogenic arrest starting from step 9 of spermiogenesis with spermatid phagocytosis and degradation. Co-immunoprecipitation and mass spec analyses revealed an interaction of ACTL7B with dynein light chains DYNLL1 and DYNLL2. Loss of ACTL7B leads to mislocalization of DYNLL1 and DYNLL2 in spermatids of *Actl7b-/-* males highlighting its role in cytoskeletal re-organization.

## Results

### Generation of *Actl7b*-deficient mice

We applied CRSIPR/Cas9-mediated gene editing in oocytes to generate *Actl7b*-deficient mice. Two guides were used targeting the intron-less coding sequence of *Actl7b* (**Fig. 1A**, arrowheads). Founders were backcrossed to C57BL/6J mice and the *Actl7b* locus was sequenced in the F1 generation. A mouse carrying a 473 bp deletion causing a frameshift was selected to establish an *Actl7b-*deficient line and animals were analyzed starting from generation N2. A genotyping PCR was established to discriminate the *Actl7bΔ* from the WT allele (**Fig. 1B**). *Actl7b* was described to be expressed in round and elongating spermatids in mice and human (Hisano et al., 2003, Guo et al., 2018, Lukassen et al., 2018) (**Fig. 1C, Fig. S1**).

**Fig. 1.**
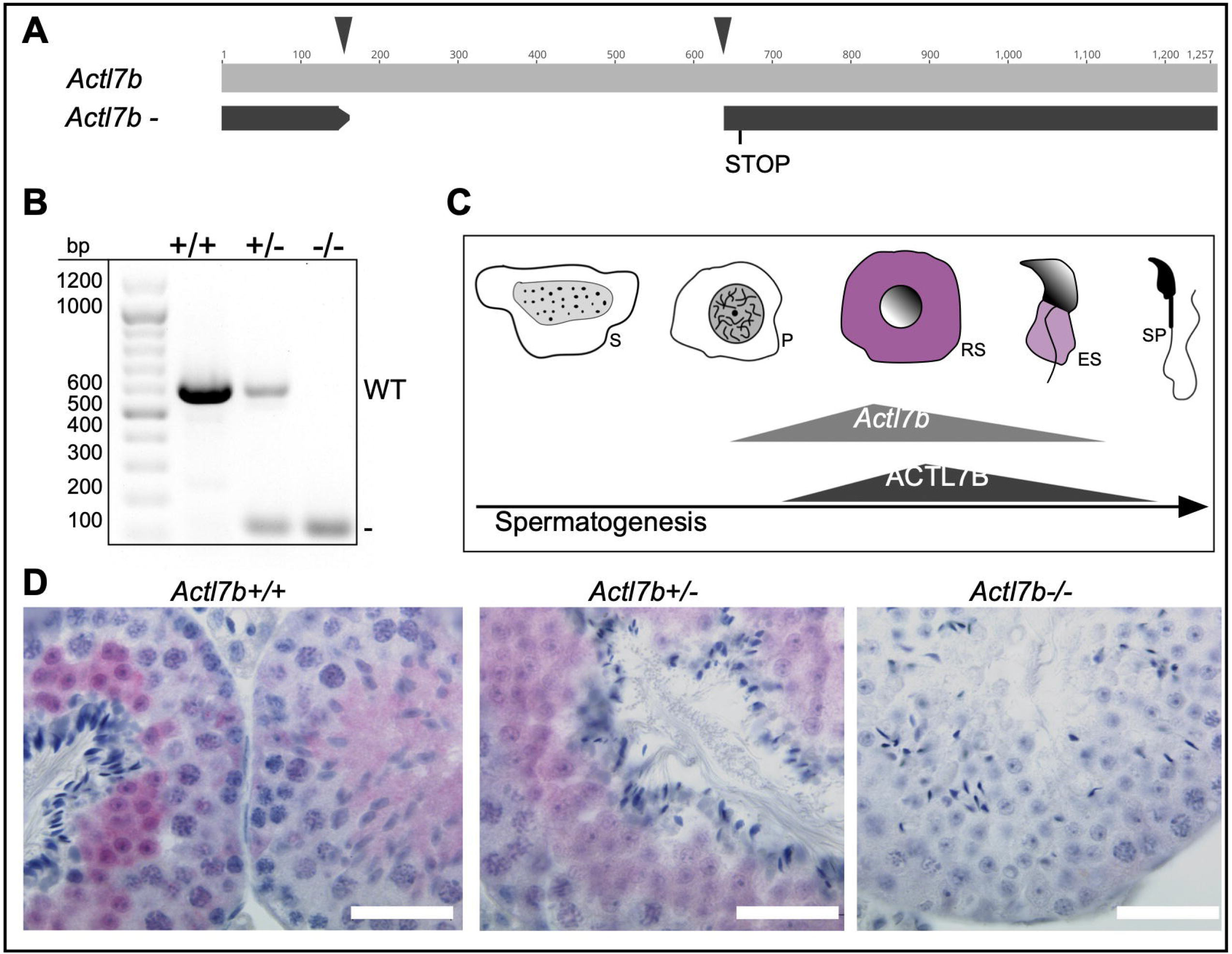
Establishment of *Actl7b*-deficient mice. **(A)** Graphical representation of CRISPR-Cas9-mediated gene editing of the *Actl7b* locus using two guide RNAs (indicated by black arrow heads) targeting the intron-less *Actl7b* coding sequence. 473 bp were deleted causing a frameshift leading to a premature stop. **(B)** Agarose gel of genotyping polymerase chain reaction of *Actl7b+/+, Actl7b+/-* and *Actl7b-/-* mice (WT band: 607 bp; KO band:134 bp). bp= base pairs **(C)** Graphical representation of *Actl7b* expression and ACTL7B immunolocalization during spermiogenesis based on literature (Hisano *et al*. 2003b, Tanaka *et al*. 2003, Guo *et al*. 2018). **(D)** Immunohistochemical staining against ACTL7B on Bouin-fixed, paraffin-embedded *Actl7b+/+, Actl7b+/-* and *Actl7b-/-* testis sections counterstained with hematoxylin. Scale: 20 μm

We used testis sections from heterozygous (*Actl7b+/-*), homozygous (*Actl7b-/-*) and WT mice (*Actl7b+/+*) for immunohistochemical (IHC) staining against ACTL7B. We demonstrate, that ACTL7B localizes to the cytoplasm of round and elongating spermatids (**Fig. 1D**), in WT and heterozygous animals confirming published data (Tanaka et al., 2003). Of note, the staining is intense in round spermatids and weakens as spermatids elongate. In testis sections of *Actl7b+/-* mice, ACTL7B signal appeared weaker suggestive of a gene-dosage effect. No staining was detected in *Actl7b-/-* testis sections, validating the null-allele.

### *Actl7b-*deficiency leads to a disruption of spermatogenesis and infertility in male mice

Fertility analysis revealed that *Actl7b+/-* males produce similar litter sizes and pregnancy frequencies as *Actl7b+/+* males (**Fig. 2A-B**), while *Actl7b-/-* males are infertile. Macroscopic analysis of the reproductive organs shows that testis weight, testis to body weight ratio and testis size of *Actl7b-/-* males is significantly reduced compared to *Actl7b+/-* and *Actl7b+/+* males (**Fig. 2C-E**), indicating defects in spermatogenesis. Next, histological sections of testis, caput and cauda epididymis were prepared. Sperm production appears normal in *Actl7b+/-* males (**Fig. 2F**). *Actl7b+/-* testis sections containing step 13-15 spermatids do not significantly differ from WT testis sections and epididymides contain mature, morphologically normal sperm. In contrast, spermatogenesis in *Actl7b-/-* males appears to be disrupted. Caput and cauda epididymides are filled with cell debris, morphologically abnormal spermatids and roundish cells most likely represent immature germ cells (**Fig. 2F**).

**Fig. 2.**
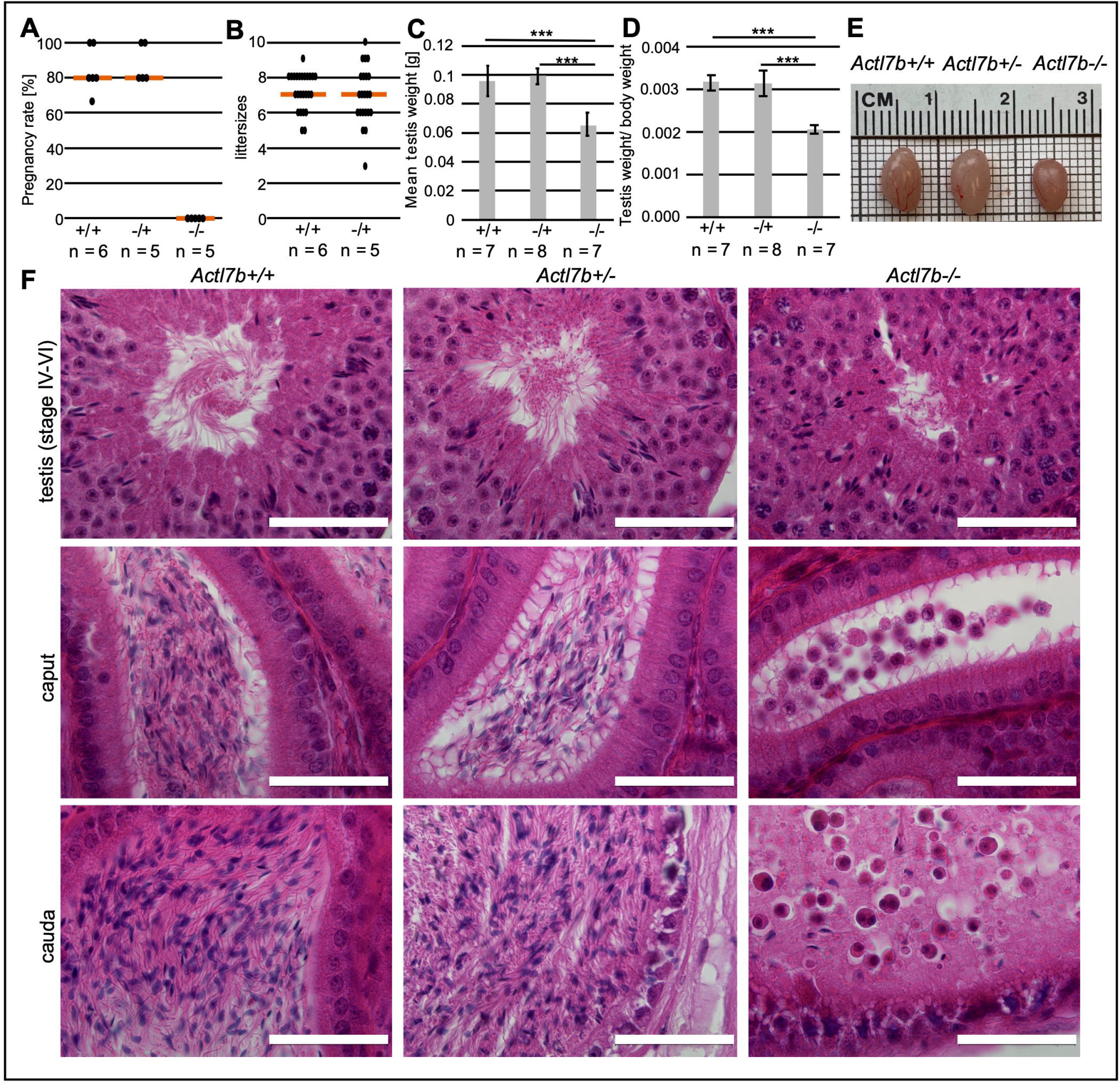
Fertility analysis and reproductive organ morphology. **(A)** Pregnancy rate of *Actl7b+/+, Actl7b+/-* and *Actl7b-/-* males mated with female WT C57BL/6J mice (n = number of males). **(B)** Average litter sizes monitored after mating of *Actl7b+/+, Actl7b+/-* males with female WT C57BL/6J mice (n = number of males). 5 plugs per male were monitored. **(C)** Mean testis weight of *Actl7b+/+, Actl7b+/-* and *Actl7b-/-* males (n = number of males). **(D)** Testis to body weight ratio of *Actl7b+/+, Actl7b+/-* and *Actl7b-/-* males (n = number of males). **(E)** Representative picture of testis dissected from *Actl7b+/+, Actl7b+/-* and *Actl7b-/-* littermates with similar body weight. **(F)** Hematoxylin-Eosin staining of testis (IV-VI of the epithelial cycle), caput epididymis and cauda epididymis of *Actl7b+/+, Actl7b+/-* and *Actl7b-/-* males. Scale: 50 μm

*Actl7b-/-* seminiferous tubules appear disorganized and germ cell development abnormal (**Fig. 3A-B**). Vacuolations in the seminiferous tubules are detected mostly in the basal region of the seminiferous tubules, indicating recent loss of germ cells. In stage VIII, retained abnormally formed elongated spermatids originating from the previous cycle can be seen. Importnatly, we detected round spermatids that seem to be blocked in development and present with dark cytoplasm, indicative for apoptosis/degradation. Fully developed, morphologically normal step 16 spermatozoa are not detected. Immature germ cells seem to be released into the lumen of seminiferous tubules. Late stage spermatids show abnormal morphologies. Various developmental stages, which usually are not found together in the same seminiferous tubule can be detected (**Fig. 3B**). Flagellar structures are detected in clusters throughout the whole tissue, even close to the basal lamina. In *Actl7b-/-* seminiferous tubules, vesicles filled with degrading spermatids were detected (**Fig. 3C-D**). These contained condensed nuclei, acrosomal and tail structures, mitochondria and granular material. This suggests, that Sertoli cells have engulfed and are degrading apoptotic spermatids.

**Fig. 3.**
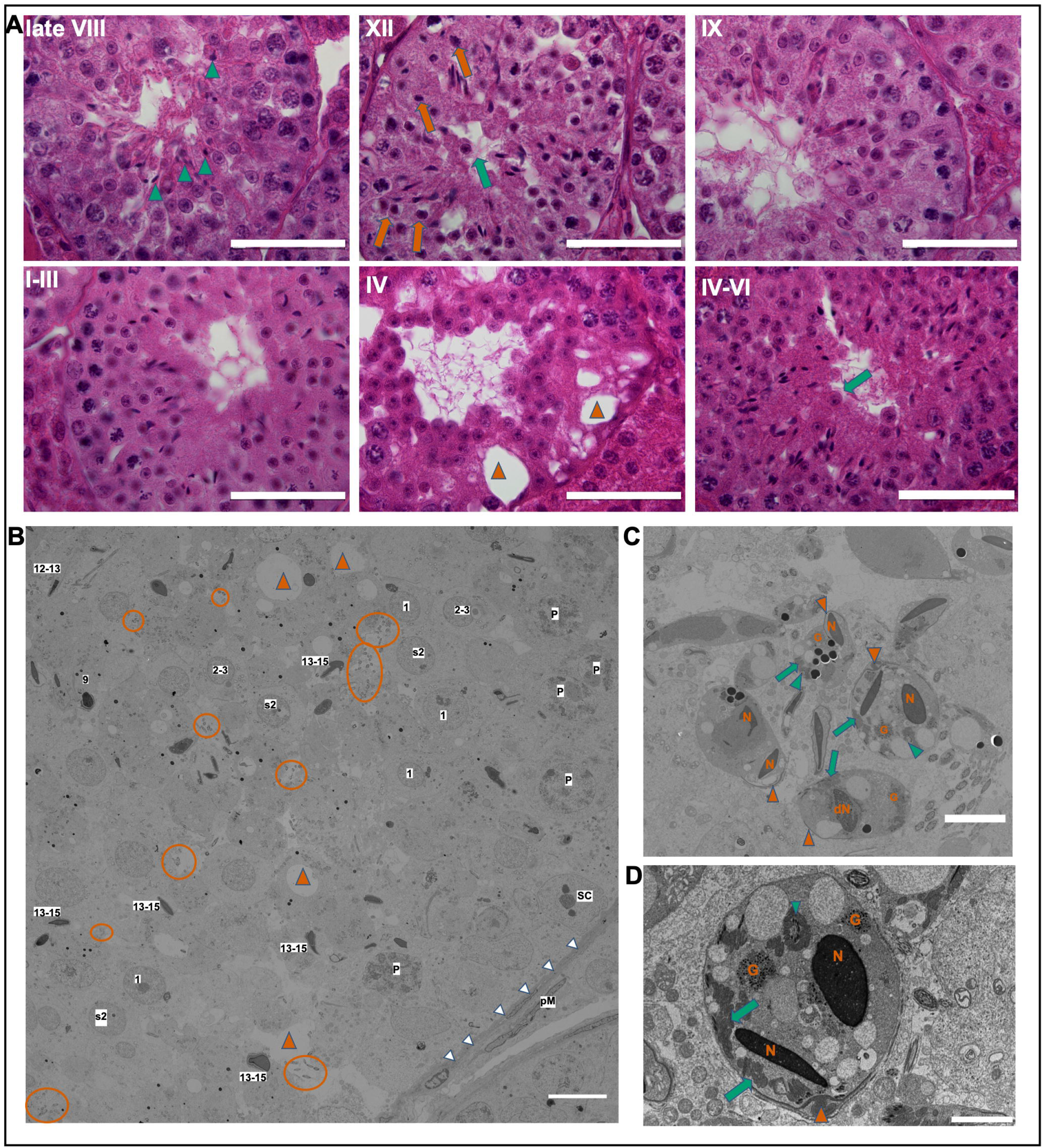
Morphology of *Actl7b*-deficient seminiferous tubules. **(A)** Hematoxylin-Eosin staining of Bouin-fixed paraffin-embedded testis sections of *Actl7b-/-* mice. Immature apoptotic germ cells can be seen to be released into the lumen (marked by green arrows). In late stage VIII elongated spermatids with an abnormal morphology, which were not spermiated, were seen (marked by green arrow heads) and round spermatids blocked in development with dark cytoplasm were found (marked by vermillion arrows). Vacuolation of seminiferous tubules was detected (marked by vermillion arrow heads). Scale: 50 µm **(B)** Representative transmission electron micrograph of a disorganized Actl7b-/- seminiferous tubule. vermillion arrow heads = vacuoles, encircled in vermillion = flagellar cross sections, s2 = secondary spermatocyte, 1 = step 1 spermatid with centrally positioned nucleus and chromatoid body, 2-3 = step 2-3 spermatid with centrally positioned nucleus and acrosomal vesicle not yet connected to nucleus, 9 = step 9 spermatid with approaching centriole, 12-13 = step 12-13 spermatid with not yet fully condensed chromatin and manchette, 13-15 = abnormal step 13-15 spermatids, eP = early pachytene spermatocytes, SC = Sertoli cell nucleus, white arrow heads = basal lamina, pM = peritubular myoid cell, Scale: 10 µm **(C-D)** Transmission electron micrographs of vesicles filled with degrading spermatids detected in Actl7b-/- seminiferous tubules. N = condensed nuclei, dN = degraded nucleus, G = granular material, vermillion arrow heads = acrosomal structures, green arrow heads = flagellar cross sections, green arrows = mitochondria. Scales: 5 µm, 2 µm

While *Actl7b-/-* males show a pathomorphological phenotype, loss of one allele of *Actl7b* seems to be phenotypically inconspicuous. We isolated sperm from the cauda epididymides of *Actl7b+/-* and WT males (**Fig. S2**). Sperm count and the percentage of viable sperm is not significantly different between *Actl7b+/-* and WT males. *Actl7b+/-* sperm appear morphologically normal, viable and motile.

### Spermiogenesis is disrupted in *Actl7b-/-* males and *Actl7b-/-* spermatids show various structural defects

To examine spermiogenesis in *Actl7b*-deficient males in more detail, we looked at basic parameters including acrosome biogenesis, DNA condensation and sperm tail formation. Since, ACTL7B is first present in round spermatids, effects were expected to manifest from round spermatid stage onwards. In the *Actl7b-/-* testis spermatogenesis arrests later at elongating spermatid stages (**Fig. 4A**). In comparison to *Actl7b+/-* and *Actl7b+/+*, in *Actl7b-/-* testis, acrosomal structures are less frequent and disorganized after spermatids start to elongate. Generally, less spermatids develop in *Actl7b-/-* testis. A positive signal for ODF2 in the lumen of seminiferous tubules of *Actl7b-/-, Actl7b+/-* and *Actl7b+/+* testis sections indicated that sperm flagellar structures are formed (**Fig. 4B**). Staining against PRM2 in the nuclei of elongating spermatids in *Actl7b-/-, Actl7b+/-* and *Actl7b+/+* testis sections indicate that nuclear remodeling and chromatin condensation is initiated. However, staining against ODF2 and PRM2 in *Actl7b-/-* seminiferous tubules appears more dispersed due to the impairment in spermatogenesis.

**Fig. 4.**
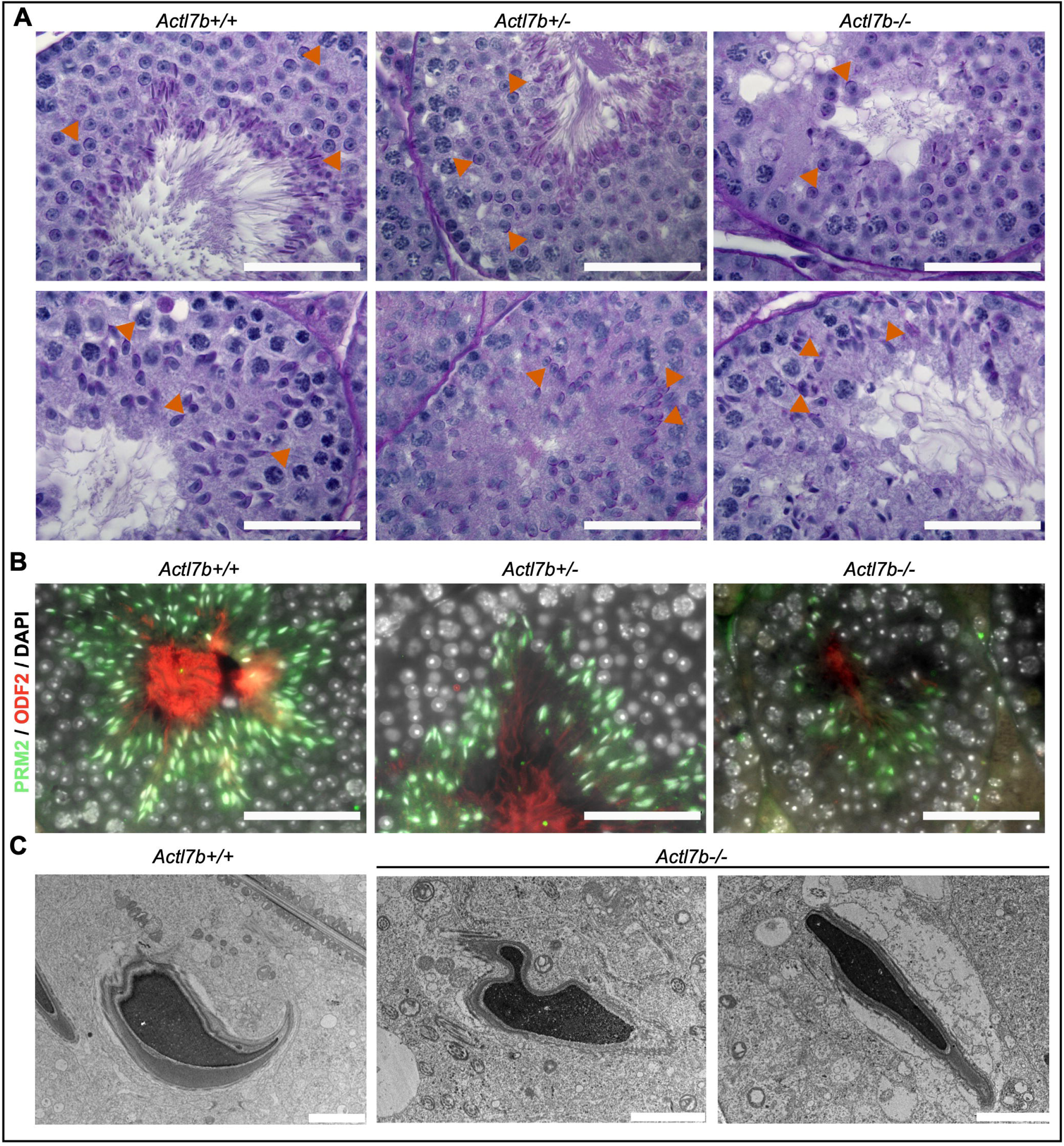
Acrosome formation, flagella formation and chromatin condensation in *Actl7b*-deficient mice. **(A)** Periodic acid Schiff staining of testis of *Actl7b+/+, Actl7b+/-* and *Actl7b-/-* males. Acrosomal structures are indicated by vermillion arrow heads. Scale: 50 μm **(B)** IHC staining against ODF2 (red fluorescence) and PRM2 (green fluorescence) of *Actl7b+/+, Actl7b+/-* and *Actl7b-/-* testis tissue sections. DAPI (in grey) was used as the counterstain. Scale: 50 μm **(C)** Transmission electron micrographs of testicular sperm presented with condensed chromatin from *Actl7b+/+* and *Actl7b-/-* males. Scale: 2 µm

In transmission electron micrographs of the testis some spermatid heads with electron dense nuclei characteristic for condensed DNA were detected in *Actl7b-/-* testis (**Fig. 4C**). In these few cells, electron density appears similar in *Actl7b-/-* and *Actl7b+/+* spermatids. Closer inspection of transmission electron micrographs reveals that spermatids in *Actl7b-/-* testes show abnormal morphologies and excess cytoplasm indicative of defective eviction of cytoplasm (**Fig. 5A**). In comparison, in the lumen of *Actl7b+/+* tubules morphologically normal sperm line up to be spermiated (**Fig. 5B**). In later developmental stages in *Actl7b-/-* spermatids the sperm membranes and acrosomal structures become detached (**Fig. 5C-D**). Part of the condensed nuclei show inclusions (**Fig. 5E**) and the general architecture of elongating spermatids appears disrupted (**Fig. 5F**). Sperm specific structures fail to assemble correctly. When analyzing the single steps of spermiogenesis first defects become apparent at step 9 (**Fig. S3, Fig. S4**). A subset of spermatids lacks the manchette, several spermatids show partly detached ectoplasmic F-Actin bundles. However, none of these defects can be found consistently in all step 9 spermatids. Defects become more pronounced in spermatids in later steps of spermiogenesis (**Fig. S4**). In part of the developing spermatids, chromatin condensation seems to be initiated earlier compared to WT spermatids. Darkly stained chromatin could, however, also be a sign of DNA degradation. Irregular sperm head shaping becomes apparent already in step 9 spermatids. The proximal centriole locates to the posterior region of the nucleus to form the basal body, but often the extension of the axoneme fails in *Actl7b-/-* spermatids. In later stage spermatids the flagellum is absent. However, cross-sections of properly formed flagella can be found detached from spermatids (**Fig. 5A**).

**Fig. 5.**
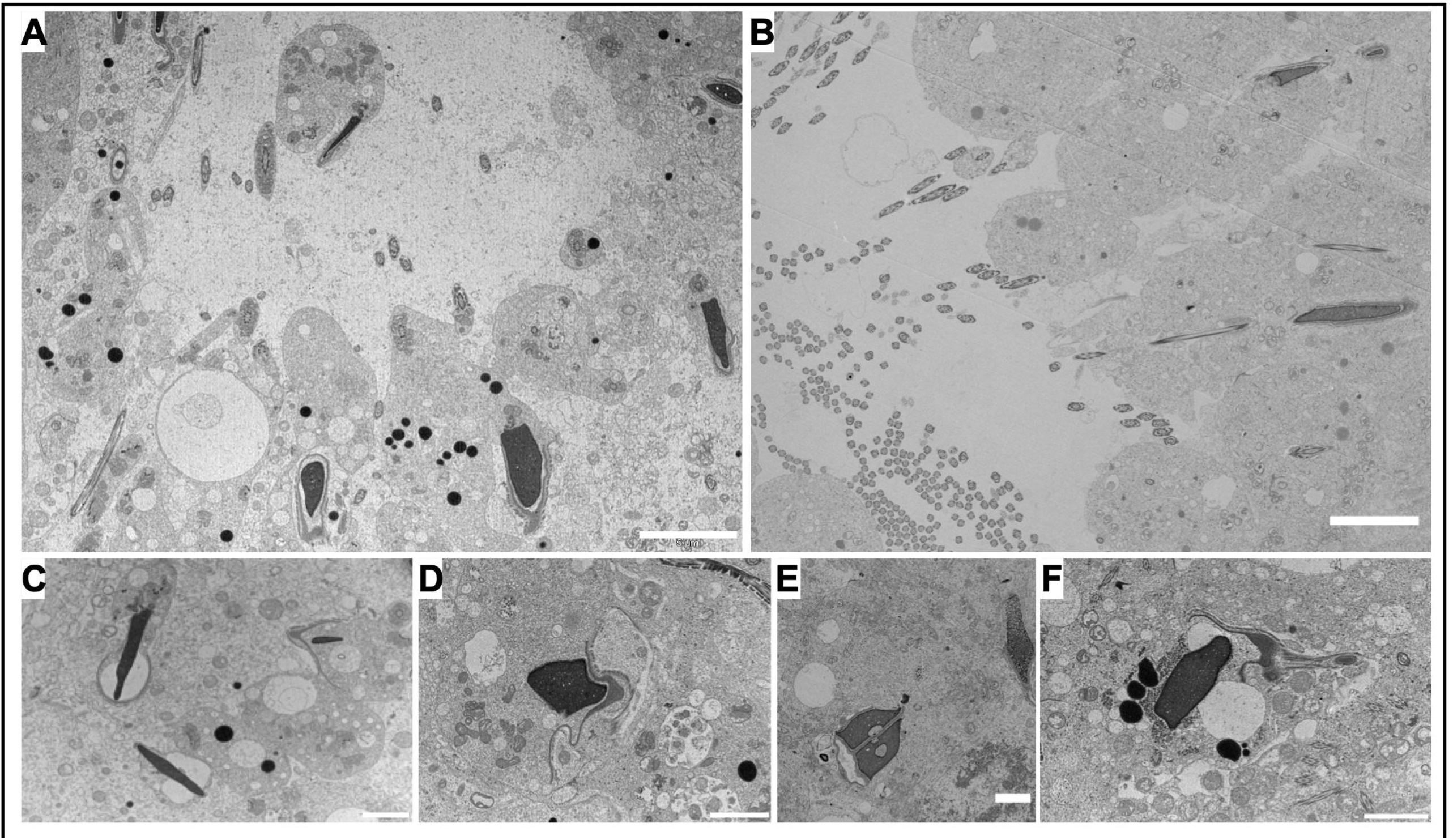
Ultrastructural analysis of *Actl7b*-/- testis. **(A)** Transmission micrograph of a lumen of an *Actl7b-/-* seminiferous tubule. **(B)** Transmission micrograph of a lumen of an *Actl7b+/+* seminiferous tubule. **(C-F)** Representative images of *Actl7b-/-* spermatids with condensed nuclei. Scales A-B: 5 µm, C-F: 2µm

These results lead us to conclude that spermiogenesis is disrupted in *Actl7b-/-* males. However, the block in development seems to be heterogeneous. While some germ cells arrest in development at round spermatid stage and appear to enter apoptosis or are released into the lumen, others develop further, form more advanced acrosomal structures, show hypercondensed chromatin and flagella formation. Apparently, key processes of sperm development are initiated, but seemingly run in an uncoordinated fashion. Finally, abnormally formed spermatids become engulfed and degraded.

### ACTL7B interacts with dynein light chains DYNLL1 and DYNLL2

To identify ACTL7B-protein interactions, anti-ACTL7B antibody was coupled to Dynabeads and used for co-immunoprecipitation (Co-IP) on protein extracts from whole WT testes. Uncoupled beads were used as control. MassSpec with the eluate was performed for protein identification (**Supplementary Material 1**). After excluding contaminating peptides like keratins, peptides identified in the “beads only” control were subtracted from the dataset. Further, a published bead proteome from HeLa cells was used to filter out proteins which nonspecifically bind Dynabeads (Trinkle-Mulcahy et al., 2008). In the Co-IP using the anti-ACTL7B-coupled beads, we identified LC8 light chains, Dynein light chain 1 (DYNLL1) and its paralog dynein light chain 2 (DYNLL2) in a similar abundance as ACTL7B.

Using western blots ACTL7B, DYNLL1 and DYNLL2 were identified in the protein input, as well as in the non-bound flow through fraction of both the Co-IP with the anti-ACTL7B-coupled beads and the Co-IP using uncoupled beads (**Fig. 6A, Fig. S5A-B**). All three proteins were detected in the eluate of the Co-IP with the anti-ACTL7B-coupled beads, but not in the eluate of the Co-IP using uncoupled beads. In the input ACTL7B is presented as a band of around 49 kDa. In the eluate of the Co-IP with the anti-ACTL7B-coupled beads, the ACLT7B band is shifted to around 60 kDa, suggesting that ACTL7B was still bound to DYNLL1 and DYNLL2. DYNLL1 and DYNLL2 present as a band of approx. 12 kDa in the input. In the eluate of the Co-IP with the anti-ACTL7B-coupled beads however, DYNLL1 and DYNLL2 present as a band of around 60 kDa, again suggesting that they are bound to ACTL7B. DYNLL2 shows weak bands at the same height already in the input. Next, Co-IP was repeated using anti-DYNLL1 and anti-DYNLL2-coupled beads (**Fig. 6B, Fig. S5C**). The anti-DYNLL1 antibody appeared not to be suitable for coupling and/or Co-IP, since no DYNLL1 was detected in the eluate (not shown). In the eluate of the Co-IP using anti-DYNLL2-coupled beads, DYNLL2 and ACTL7B were detected, validating ACTL7B-DYNLL2 interaction. Western Blots on protein extraction from *Actl7b-/-, Actl7b+/-* and *Actl7b+/+* testis, showed that DYNLL1 and DYNLL2 protein amounts are not reduced in *Actl7b-/-* testis compared to *Actl7b+/-* and *Actl7b+/+* testes (**Fig. 6C**).

**Fig. 6.**
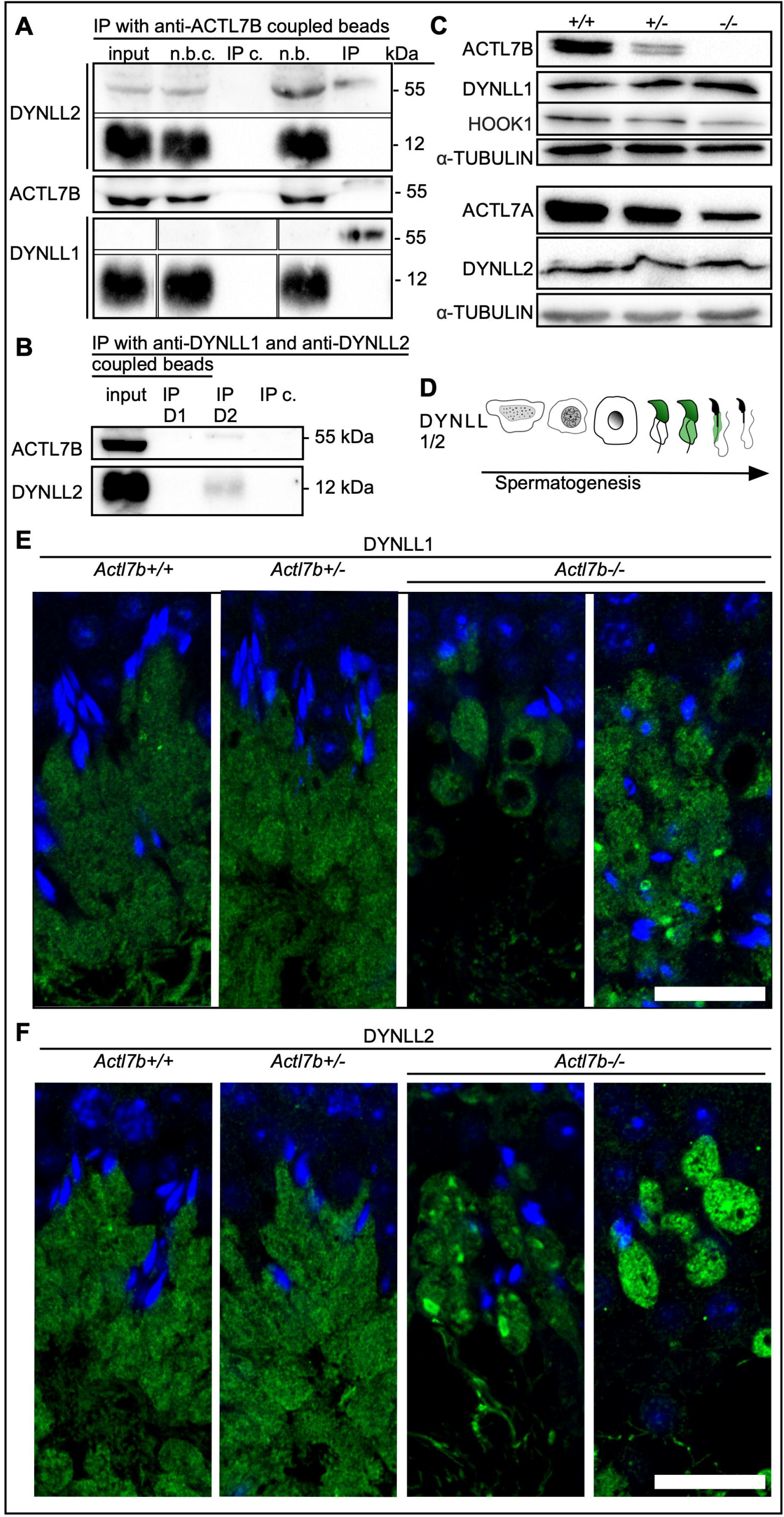
Interaction of ACTL7B with DYNLL1 and DYNLL2. **(A)** Western blots on the protein input (whole WT testis), the non-bound fraction of the beads only control (n.b. c.), the IP eluate of the beads only control (IP c.), non-bound fraction of the anti-ACTL7B-coupled beads (n.b.), the IP eluate of the anti-ACTL7B-coupled beads (IP). Anti-DYNLL2, anti-DYNLL1 and anti-ACTL7B were used. **(B)** Western blots on the protein input (whole WT testis), the IP eluate of the anti-DYNLL1-coupled beads (IP D1), the IP eluate of the anti-DYNLL2-coupled beads (IP D2), the IP eluate of the beads only control (IP c.). Anti-ACTL7B and anti-DYNLL2 were used for western blots. **(C)** Western blots on protein extractions from whole *Actl7b+/+, Actl7b+/-* and *Actl7b-/-* testis. Anti-ACTL7B, anti-DYNLL2, anti-HOOK1, anti-ACTL7A and anti-DYNLL2 were used. α-tubulin was used as loading control. **(D)** Graphical depiction of DYNLL1 and DYNLL2 immunolocalization during spermiogenesis based on literature (Wang et al., 2005). **(E)** DYNLL1 staining in *Actl7b+/+, Actl7b+/-* and *Actl7b-/-* elongating spermatids. Dapi was used as counterstain. Scale: 20 µm. **(F)** DYNLL2 staining in *Actl7b+/+, Actl7b+/-* and *Actl7b-/-* elongating spermatids. Dapi was used as counterstain. Scale: 20 µm.

To evaluate potential compensation of ACTL7B by ACTL7A, ACTL7A abundance was analyzed by western blot. ACTL7A levels are reduced in *Actl7b-/-* testes compared to *Actl7b+/-* and *Actl7b+/+* testes, suggesting no compensatory function of ACTL7A. Further, HOOK1, a protein expressed late in spermiogenesis, was also reduced in *Actl7b-/-* testis. We hypothesize that reduced protein level expressed late in spermiogenesis (like HOOK1) is due to the reduced numbers of spermatids reaching later stages of spermiogenesis in *Actl7b-/-* mice.

DYNLL1 is described to be localized first to the nucleus of elongating spermatids and later in the cytoplasm and residual bodies (**Fig. 6D**) (Wang et al., 2005). Immunofluorescent stainings against DYNLL1 and DYNLL2 revealed that both proteins show the identical localization. As described, they localize to the head of early elongating spermatids and were present in the cytoplasm at later developmental stages (**Fig. S6, Fig. S7**). Interestingly, DYNLL1 and DYNLL2 expression correlates with the onset of defects observed in spermatids of *Actl7b-/-* mice. In the *Actl7b-/-* testis, the first clear differences in the DYNLL1 and DYNLL2 stainings compared to WT were seen at around stage I-III of the seminiferous cycle. Both light chains should be homogeneously present throughout the whole cytoplasm, which should be orientated towards the lumen in a stream-like fashion. In *Actl7b-/-* testes, however, the DYNLL1/2-positive cytoplasm is arranged in roundish sacs (**Fig. 6E-F, Fig. S6, Fig. S7**). Vacuolation of the staining and foci of concentrated DYNLL1/2 were detected. Taken together these results suggest, that the localization of DYNLL1 and DYNLL2 is altered in the absence of ACTL7B, while the amount of protein is unchanged. Of note, F-/G-actin ratios are not significantly different in *Actl7b-/-* compared to *Actl7b+/+* or *Actl7b+/-* testis, suggesting that actin filament turnover is unlikely to be affected in the absence of ACTL7B (**Fig. S8A**). Even though the Co-IP did not reveal ACTL7B-actin interaction, actin filament arrangement is disturbed in some areas of *Actl7b*-/- seminiferous tubules (**Fig. S8B**).

### Loss of ACTL7B leads to proteomic changes in *Actl7b-/-* testis

In order to analyze alterations in the testicular proteome in *Actl7b*-deficient mice, protein samples isolated from whole testes of each five *Actl7b-/-, Actl7b+/-* and *Aclt7b+/+* mice were used for mass spectrometric analysis (**Supplementary material 2**). Principal component analysis (PCA) showed that the *Actl7b-/-* samples cluster apart from the *Actl7b+/-* and *Actl7b+/+* samples (**Fig. S9A**). Differential abundance (DA) analysis revealed no significant difference in protein abundance in *Actl7b+/-* compared to *Actl7b+/+* samples (**Fig. 7A, Fig. S9B**). Differentially abundant proteins were detected in *Actl7b-/-* samples compared to *Actl7b+/-* and *Actl7b+/+* samples, respectively (**Fig. 7B-C**). 30 proteins were higher abundant and 9 proteins lower abundant in *Actl7b-/-* compared to *Actl7b+/+* samples and 24 proteins were higher abundant and 10 proteins lower abundant in *Actl7b-/-* compared to *Actl7b+/-* with a stringent log fold change (LFC) of ≥1 (**Fig. 7D-E, Fig. S9B**). 19 of the higher abundant proteins were detected in both comparisons (**Fig. 7D**).

**Fig. 7.**
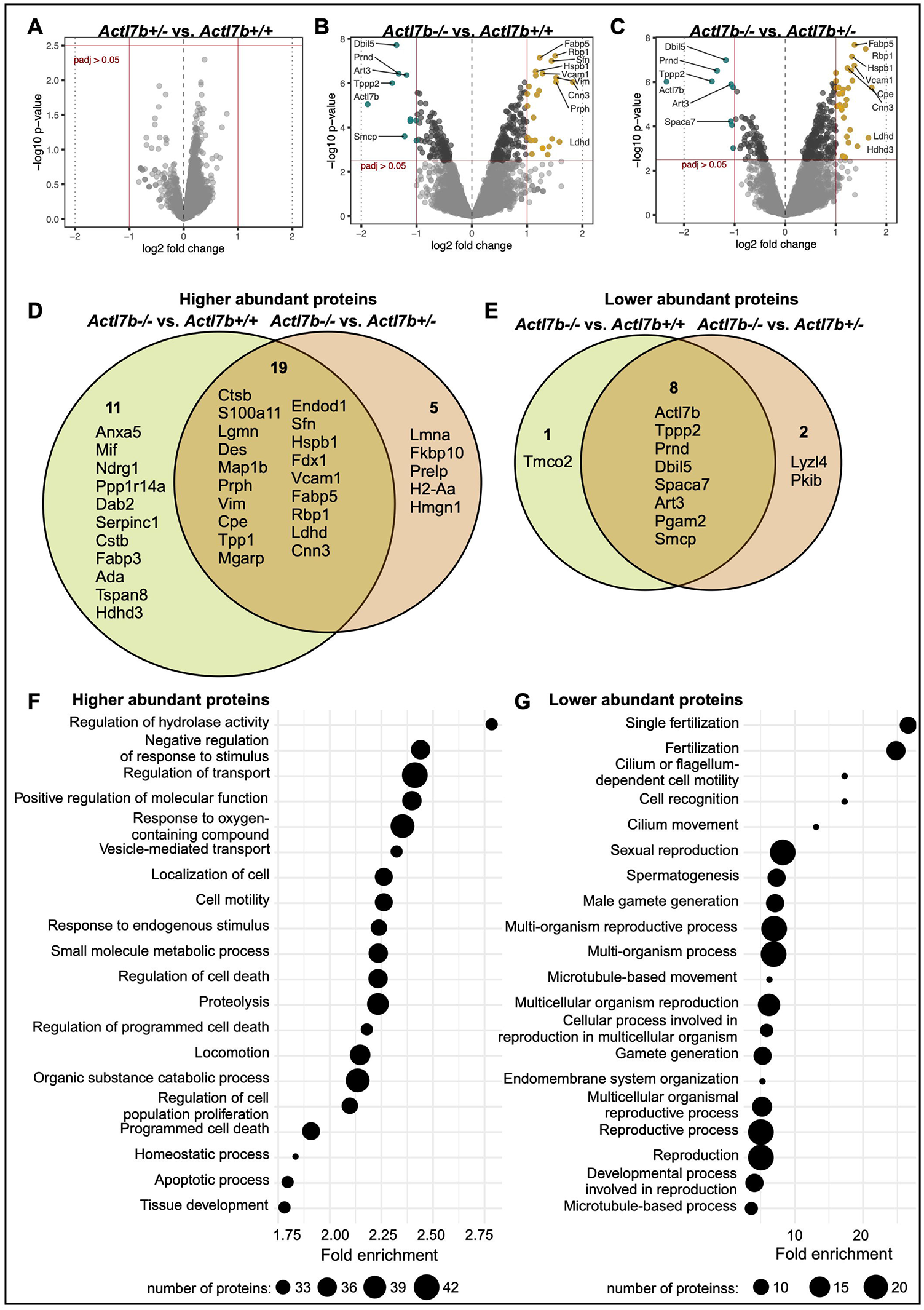
Proteomic analysis of *Actl7b*-deficient testis. **(A-C)** Volcano plots showing differential abundance (DA) of proteins in *Actl7b+/-* compared to *Actl7b+/+* **(A)**, *Actl7b-/-* compared to *Actl7b+/-* **(B)** and *Actl7b-/-* compared to *Actl7b+/+* **(C)** testis. Proteins showing a significant DA are indicated in teal (lower abundant) and yellow (higher abundant) (adjusted p-value > 0.05). Top DA proteins are labeled with their corresponding gene symbol. **(D-E)** Venn diagrams showing the overlap of significantly higher **(D)** and lower **(E)** abundant proteins in the comparisons of *Actl7b-/-* vs. *Actl7b+/+* and *Actl7b-/-* vs. *Actl7b+/+* testis (adjusted p-value ≤ 0.05, LFC ≤ 1). **(F-G)** Dot plots of the top twenty gene ontology (GO)-biological process terms significantly enriched in the higher abundant **(F)** and lower abundant **(G)** proteins in *Actl7b-/-* vs. *Actl7b+/+* samples (p ≤ 0.05, LFC ≤ 0.05). Dot sizes represent the number of proteins contributing to the GO-term.

Several Sertoli cell expressed proteins, such as the type III intermediate filament proteins vimentin, desmin and peripherin (Mruk and Cheng, 2004, Vogl et al., 2008) as well as fatty acid-binding protein 5 (FABP5) and intracellular retinol-binding protein 1 (RBP1) (Oresti et al., 2013, Griswold, 2022), were found to be higher abundant in *Actl7b-/-* testicular samples. Differential abundance of these mostly Sertoli cell enriched proteins might indicate a secondary effect caused by arrested and apoptotic germ cells. Moreover, proteins associated with protein or nucleic acid degradation as well as early apoptosis were higher abundant in *Actl7b-/-* testis (ANXA5, CTSB, LGMN, TPP1, ENDOD1). Further, VCAM-1, proposed to function as an adhesion protein in immunoregulation of the testis was significantly higher abundant in *Actl7b-/-* testis (Sainio-Pollanen et al., 1997). As expected, ACTL7B was detected among the significantly lower abundant proteins in *Actl7b-/-* compared to *Actl7b+/-* and *Actl7b+/+* samples. Additional spermatocyte and spermatid related proteins were detected to be lower abundant in *Actl7b-/-* samples (ART3, SPACA7, SMCP, TPPP2, TMCO2, PGAM2, LYZL4, PRND, PKIB).

When applying a less stringent LFC of ≥0.5, 193 proteins were higher abundant and 59 proteins lower abundant in *Actl7b-/-* compared to *Actl7b+/+* samples (**Fig. S9B, Tab. S1**). These proteins were used to analyze the enrichment in biological processes (**Fig. 7F-G**). Proteins higher abundant in *Actl7b-/-* compared to *Actl7b+/+* showed an enrichment in protein degradation processes, apoptosis, oxidative stress response, cell motility and localization (**Fig. 7F**). Interestingly, N-cadherin (gene symbol: *Cdh2*) was higher abundant in *Actl7b-/-* samples. In Sertoli cells N-cadherin is required for blood-testis-barrier integrity (Jiang et al., 2015). Sertoli cell N-cadherin interacts with actin and vimentin and is found at the ectoplasmic specializations between Sertoli cells and germ cells and in the basal compartment of the seminiferous tubules (Mruk and Cheng, 2004). Ezrin, which was also higher abundant in *Actl7b-/-* testis, regulates Sertoli cell-spermatid-adhesion, influences spermatid polarity and is involved in residual body/phagosome transport (Gungor-Ordueri et al., 2014). It has been shown that in cells transfected with a mutated ezrin, tubulin, actin and vimentin structure is altered (Zhang et al., 2020). Furthermore, it has been shown that ezrin interacts with VCAM-1 *in vitro* (Barreiro et al., 2002). IHC staining against ezrin showed an intense accumulation of ezrin around germ cells and vacuolations in *Actl7b-/-* seminiferous tubules, indicating increased germ cell transport and clearance (**Fig. S10**).

On the other hand, lower abundant proteins, on the other hand, showed an enrichment in male gamete formation, fertilization and reproduction, as well as microtubule-based movement (**Fig. 7G**).

### *ACTL7A* and *ACTL7B* are highly conserved across primates and rodents

Since *ACTL7A* and *ACTL7B* are both testis specific and show sequence similarity, we performed evolutionary analysis of both genes to compare their levels of sequence conservation and predict their essentiality. *ACTL7A* and *ACTL7B* show 57% amino acid and 52% coding sequence identity in *Mus musculus*. Of note, human and mouse ACTL7B show 85.9% pairwise amino acid identity. Selective pressures on the codon level were assessed via the nonsynonymous/synonymous substitution rate ratio (ω = dN/dS). This ratio distinguishes between purifying selection (codon sequence conservation) (ω<1), neutral evolution (ω=1) and positive selection (ω>1). Analysis of the selective constrains on *ACTL7A* and *ACTL7B* revealed that both are under strong purifying selection across primates and rodents (**Tab. 1, Fig. S11**). The evolutionary rates (ω) of the whole sequences across all included species trees were significantly lower than 1 (*ACTL7A*: ω = 0.17, p > 0.001; *ACTL7B*: ω = 0.06, p > 0.001), with *ACTL7B* showing an even lower evolutionary rate compared to *ACTL7A*. When comparing selective pressures between primates and rodents we found no significant difference in evolutionary rate between the clades. Hence, *ACTL7A* and *ACTL7B* are equally highly conserved in both clades. Lastly, the selective constraints acting on each codon site were calculated across the whole tree. 97% of the *ACTL7B* codon sites were conserved, further confirming that *ACTL7B* is under strong purifying selection. Similarly, 83% of the codon sites were conserved for *ACTL7A*. These results indicate that changes in the coding sequences, such as mutations are highly detrimental and are strongly selected against. This usually indicates that the gene and its protein product are highly essential. *ACTL7B* seems to be more strongly conserved than *ACTL7A*. Additionally, since both rodents and primate clades, including the murine and human sequence are equally conserved, we can surmise that *ACTL7B* is highly essential for both human and mouse spermatogenesis.

**Table 1:** Evolutionary analysis of *ACTL7A* and *ACTL7B*. Values of selective constrains acting on *ACTL7A* and *ACTL7B* across the whole phylogenetic tree, on the clades primates and rodentia, and the codon sites across the whole tree. Results of the evolutionary analysis using CodeML (PAML4.9).

| Selection on: | | LnL null model | LnL alternative model | LRT | p | $\omega$ | interpretation | | | | | |
| --- | --- | --- | --- | --- | --- | --- | --- | --- | --- | --- | --- | --- |
| whole sequence, whole tree | | 12123.91 (M0fix) | 12510.29 (M0) | -772.77 | >0.001 | 0.17 (M0) | $\omega$ sign. different from 1 | overall conserved | | | | |
| whole sequence, clades / lineages | Primates | 12510.29 (M0) | 12507.93 (MCfree) | 4.72 | n.s. | 0.20 (M0) | $\omega$ not group specific | both lineages equally conserved | | | | |
|  | Rodentia |  |  |  |  | 0.16 (M0) |  |  |  |  |  |  |
| codon sites, whole tree |  | 12281.27 (M1a) | 12281.27 (M2a) | 0.00 | n.s. | n.a. | purifying selection on sites | sequence is 83% conserved | prop 0 | prop 1 | prop 2a | PSS |
|  |  |  |  |  |  |  |  |  | 0.83 | 0.17 | n.a. | n.a. |

***ACTL7B***
| Selection on: | | LnL null model | LnL alternative model | LRT | p | $\omega$ | interpretation | | | | | |
| --- | --- | --- | --- | --- | --- | --- | --- | --- | --- | --- | --- | --- |
| whole sequence, whole tree | | 12692.61 (M0fix) | 11332.97 (M0) | 2719.27 | >0.001 | 0.06 (M0) | $\omega$ sign. different from 1 | overall conserved | | | | |
| whole sequence, clades / lineages | Primates | 11332.97 (M0) | 11332.94 (MCfree) | 0.06 | n.s. | 0.06 (M0) | $\omega$ not group specific | both lineages equally conserved | | | | |
|  | Rodentia |  |  |  |  | 0.06 (M0) |  |  |  |  |  |  |
| codon sites, whole tree |  | 11273.60 (M1a) | 11273.60 (M2a) | 0.00 | n.s. | n.a. | purifying selection on sites | sequence is 97% conserved | prop 0 | prop 1 | prop 2a | PSS |
|  |  |  |  |  |  |  |  |  | 0.97 | 0.03 | n.a. | n.a. |

LnL = log Likelihood value of the model; LRT = Likelihood Ratio Test; p = p-value of the LRT; $\omega$ = evolutionary rate (dN/dS); sign. = significantly; prop = proportion of sites assigned to the respective site class; PSS = positively selected sites; for explanation of the models see M&M.

## Discussion

Here, we have generated an *Actl7b*-deficient mouse model to analyze the role of ACTL7B in spermatogenesis. Loss of ACTL7B led to male infertility in mice due to absence of functional, mature sperm. *Actl7b-/-* spermatids showed arrest in development resulting in a wide variety of abnormalities starting at step 9, such as detached acrosomes and membranes, failed elongation of the axoneme and impaired eviction of excess cytoplasm. Spermatids were subsequently degraded. A subset of degrading and immature spermatids was released into the lumen of the seminiferous tubules. Vesicles containing degrading spermatids seem to be eliminated by Sertoli cells. *Actl7b+/-* males showed reduced ACTL7B levels but remained fertile, producing motile, viable sperm at concentrations similar to WT. Since the genes of human and mouse *ACTL7B* are highly similar, we can surmise that ACTL7B variants can lead to failed spermatogenesis and infertility in humans.

Interestingly, the actin related protein ACTL7B does not seem to talk to the actin cytoskeleton, since we could not find a change in steady state actin filaments. Rather, ACTL7B seems to link to the microtubule network related functions.

By Mass Spec analyses we demonstrated that ACTL7B interacts with dynein light chains DYNLL1 and DYNLL2. Of note, ACTL7B becomes upregulated in round spermatids while dynein light chains DYNLL1 and DYNLL2 are first detected later, in step 9 spermatids (Wang et al., 2005). This correlates with the onset of the defects in spermatogenesis of *Actl7b-/-* males, suggesting, that ACTL7B-DYNLL1/2 interaction is crucial and leads to the defects observed. The LC8 family light chains DYNLL1 and its paralog DYNLL2 are highly conserved orthologs among mammals (Rapali et al., 2011). LC8 light chains are presumably involved in dynein complex assembly, thereby indirectly affecting cargo binding. The LC8 light chains are hub proteins that form homodimers with high conformational dynamics of binding grooves, interacting with a wide variety of proteins and function in multiple cellular processes like mitosis, intracellular transport, the stabilization of microtubules, nuclear transport, apoptosis, postsynaptic density, regulation of transcription (Rapali et al., 2011, Nyarko et al., 2011, Jespersen and Barbar, 2020, Hall et al., 2008). They bind intrinsically disordered proteins as dimers thereby linking subunits in multiprotein complexes (Reardon et al., 2020, Asthana et al., 2012). DYNLL1 was further shown to facilitate dissociation of dynactin from dynein, regulating cargo release (Jin et al., 2014). We hypothesize that DYNLL1 and DYNLL2 promote ACTL7B dimerization and stabilization in multiprotein complexes. Alternatively, ACTL7B might be involved in LC8 transport or activation, since absence of ACTL7B caused mislocalization of the LC8 light chains. Effects of the LC8 light chains on the microtubule structure or the dynein 1 motor complex might explain the phenotype seen in *Actl7b*-deficient mice.

It has been shown that knockdown of the cytoplasmic dynein 1 heavy chain DYNC1H1, causes a disruption of the microtubule structure and polymerization in Sertoli cells in rat (Wen et al., 2018). Further, F-actin organization was perturbed, spermatid polarity was affected and spermatid transport and release were defective. Lastly, phagosome transport was affected. These effects are similar to those described for *Actl7b*-deficient mice here. Another study identified SPEF2 as interaction partner of DYNC1H1 (Lehti et al., 2017). Germ cell specific knockout of *Spef2* let to multiple spermatid differentiation defects, severely reduced sperm numbers and male mice infertility. It has been proposed that SPEF2 functions as a linker protein, interacting with dynein 1 to facilitate cargo transport along microtubules during spermatid differentiation. ACTL7B might have similar functions. Other dynein light chains have been described in spermatogenesis. For the dynein light chain tctex-type 4 (DNLT4), 40 different interactors have been identified in human testis, showing the functional variety of dynein light chain interacting proteins (Freitas et al., 2014). In human, absence or lower levels of dynein light chain tctex-type 1 (DNLT1), have been associated with male infertility (Indu et al., 2015). A detailed analysis of LC8 function in murine spermatogenesis is mandatory, in order to reveal the role and consequence of ACTL7B-LC8 interactions.

Mass spectrometric analysis of testicular proteins revealed that Sertoli cell associated proteins are higher abundant in *Actl7b-/-* testis compared to *Actl7b+/-* and *Actl7b+/+* testis, indicating a reaction to defective spermatids. Intermediate filaments (vimentin, desmin and peripherin), ezrin, N-cadherin and VCAM-1 were found to be higher abundant. Intermediate filaments usually surround the Sertoli cell nucleus and from there extend to desmosome junctions, which are localized between adjacent Sertoli cells and between Sertoli cells and germ cells (Johnson, 2014). Ezrin, accumulated around germ cells and vacuolations indicating recent germ cell loss in *Actl7b-/-* testis tissue. Ezrin regulates Sertoli cell-spermatid-adhesion as well as phagosome transport (Gungor-Ordueri et al., 2014). N-cadherin localizes to ectoplasmic specializations between Sertoli cells and germ cells (Jiang et al., 2015). It interacts with intermediate filaments and actin. The vascular adhesion molecule VCAM-1, was shown to interact with ezrin *in vitro* (Barreiro et al., 2002). Under inflammation or chronic conditions, VCAM-1 expression can be activated by multiple stimuli including pro-inflammatory cytokines and ROS (Kong et al., 2018). VCAM-1 expression was been shown to be increased in Sertoli cells that have been exposed to inflammatory mediators *in vitro* (Riccioli et al., 1995). Taken together, these secondary effects of *Actl7b*-deficiency suggest rapid degradation of abnormal, developmentally blocked spermatids by Sertoli cells. Spermatid degradation correlated with a lower abundance of proteins related to spermatids, male gamete formation and fertility.

*ACTL7A* and *ACTL7B* show a high level of identity and are both expressed specifically in the testes of mice and human, which might suggest functional redundancy (Chadwick et al., 1999, Hisano et al., 2003, Tanaka et al., 2003). However, we have shown, that both genes are evolutionary conserved and under purifying selection in both rodents and primates, suggesting that both genes are required for proper sperm development and function. Further, in murine sperm ACTL7A is localized to the acrosome and tail, while ACTL7B is present in the cytoplasm of round and elongating spermatids (Fu et al., 2012, Tanaka et al., 2003). While ACTL7B is evicted and detected in residual bodies, ACTL7A is present in the acrosome of mature sperm. Together, the data strongly suggest that ACTL7A and ACTL7B have adapted to different functions. Consequently, the phenotype described for *Actl7a*-mutant mice and humans carrying Actl7a variants differ from those described here. Homozygous *ACTL7A* missense mutation causes sperm acrosomal defects and infertility in human and mice (Xin et al., 2020). *ACTL7A*-deficient sperm showed reduced levels of PLCζ, a sperm-borne oocyte activation factor. Artificial oocyte activation overcame infertility caused by *ACTL7A*-deficiency. Similar phenotypes have recently been described for human and murine sperm carrying homozygous pathogenic variants in *ACTL9* (Dai et al., 2021). Indeed, ACTL9 seems to interact with ACTL7A and both proteins are mislocalized, when ACTL9 is mutated. Another recent study identified a homozygous missense variant of *ACTL7A* in a teratozoospermic patient (Dai et al., 2022). Analysis of a mouse model carrying an equivalent mutation showed that the acrosome and acroplaxome become detached during spermiogenesis. The acroplaxome, ACTL7A and PLCζ are shed of and evicted in cytoplasmic droplets. Supporting the results of prior studies *Actl7a*-mutated sperm failed to activate the oocyte leading to infertility. All these studies showed that spermatogenesis is not disrupted when ACTL7A is missing, but acrosome formation is impaired. Hence, the phenotypes of *Actl7b* and *Actl7a*-deficiency differ greatly. Here, we additionally showed that ACTL7A levels are not elevated in *Actl7b-/-* mice, further arguing against a compensatory role of ACTL7A in *Actl7b-/-* mice.

In human, one study identified *ACTL7B* levels to be significantly lower in teratozoospermic patients compared to healthy controls (Ahn et al., 2017). Further, single nucleotide polymorphisms in *ACTL7B* have been identified in cohorts of infertile patients (Tanaka et al., 2007, Tanaka et al., 2019). These have, however, not been directly correlated to the infertility. Lastly, *ACTL7B* has been identified to convey a high discriminating power between obstructive and non-obstructive azoospermia subtypes, both on protein and transcript level, suggesting *ACTL7B* as a marker for screening patients (Davalieva et al., 2022). Our study clearly shows, that mutations in *ACTL7B* might be directly connected to male infertility, calling for further investigations in humans.

## Material and Methods

### Ethics statement

Animal experiments were performed according to the German law of animal protection and in agreement with the approval of the local institutional animal care committees Landesamt für Natur, Umwelt und Verbraucherschutz, North Rhine-Westphalia (approval ID: AZ81-0204.2018.A369).

### Generation of *Actl7b*-deficient mice

Single guide RNAs (sg1_ts: 5’-CACCCGGACACGGCGTGTCGCAT; sg1_bs: 5’-AAACCATGCGACACGCCGTGTCC; sg2_ts: 5’-CACCAATACGGAAGATCAAGGCG, sg2_bs: 5’-AAACGCGCCTTGATCTTCCGTAT) were designed using the Benchling CRISPR Guide RNA design tool (https://www.benchling.com/crispr/; ENSMUSG00000070980) and tested in ES cells as described (Schneider et al., 2016). The selected guides were ordered as crRNA sequences (IDT, Leuven, Belgium) and prepared for electroporation as described (Arévalo et al., 2022). Briefly, crRNAs were annealed to tracRNA (IDT) (50 nM) and mixed with Cas9 (IDT) in OPTI-MEM (Thermo Fisher Scientific, Waltham, USA).

CRISPR-Cas9-mediated gene editing of oocytes was performed as described before (Arévalo et al., 2022). 6-8 weeks old B6D2F1 females were superovulated by intraperitoneal injections of 5 i.u. prs serum (PMS) and 5 i.u. human chorionic gonadotropin (hCG), mated with B6D2F1 males and oocytes were isolated 0.5 dpc. Oocytes were electroporated in OPTI-MEM containing the guide RNA mix utilizing using a BioRad Gene Pulser (BioRad, Feldkirchen, Germany). After recovery and washing, the oocytes were incubated in KSOM (Merck, Darmstadt, Germany) overlaid with mineral oil at 37°C overnight.

Developing 2-cell stage embryos were transferred into the fallopian tube of pseudo-pregnant CD1 foster mice. Offspring was genotyped by PCR and sequenced to identify founder animals. Selected founders were backcrossed to C57BL/6J mice and the F1 generation was sequenced. The *Actl7bΔ* allele (NM_025271.2:c.159_631del) was further back-crossed to C57BL/6J mice. Starting from the N2 generation analyses were performed. The allele was registered with MGI (Actl7b^em1Hsc^ ; ID:671828).

### Genotyping and sequencing of mice

Primers flanking the gene edited region (Actl7b_fwd: 5’-GGGACACAGGTTCCACTCAAC, Actl7b_rev: 5’-AGGTAGTTGGTGAGGTCGCA) were used to amplify both the WT and edited allele (Cycling conditions: 5 min 95°C; 35x (30 sec 95°C; 30 sec 60°C; 45 sec 72°C); 5 min 72°C). PCR products (WT allele: 607 bp, *Actl7b-*: 134 bp) were separated on agarose gels.

Samples for sequencing were prepared as described (Merges et al., 2022) and sent to GATC/Eurofins (Cologne, Germany) for sequencing.

### Fertility assessment

Male mice, aged between 8-12 weeks, were mated 1:1/1:2 to C57BL/6J females and females were examined for presence of a vaginal plug daily. Plug positive females were separated and monitored for pregnancies and litter sizes. A minimum of five plugs per male were monitored.

### Macroscopic analysis of testis

Testis were dissected and weight was determined. Sections of Bouin-fixed testis, caput and cauda epididymis were stained with hematoxylin and eosin as described (Merges et al., 2022).

### Immunohistochemistry (IHC)/ Immunofluorescence (IF)

Tissues were fixed in Bouin’s solution (4°C, overnight) processed in paraffin and 3 µm sections were generated utilizing a microtome (Microm CP60). Heat mediated antigen retrieval was performed (pH6: anti-DYNLL1 (Invitrogen, SD08-04, 1:1500), anti-DYNLL2 (Proteintech, 16811-1-AP, 1:1500), anti-ACTL7B (Proteintech, 13537-1-AP, 1:750), anti-PRM2 (Briar Patch Biosciences, Hup2B, 1:200), anti-ODF2 (Proteintech, 12058-1-AP, 1:500), anti-Erzin (Santa Cruz, sc-58758, 1:100); pH9: anti-ACTIN (Abcam, ab179467, 1:500)). For slides stained with anti-ACTIN an additional peroxidase blocking step was performed. Slides stained against PRM2 and ODF2 were additionally treated with decondensation buffer, as described previously (Schneider et al., 2020). Sections stained with anti-DYNLL1, anti-DYNLL2 and anti-ACTIN were processed using the VectaFluor™ Horse Anti-Rabbit IgG, DyLight® 488 Antibody Kit (Vector Laboratories, Burlingame, CA, USA; DI-1788), sections stained against Ezrin were processed with the VectaFluor™ Anti-Mouse IgG, DyLight® 594 Kit (Vector Laboratories, DI-2794), sections stained against PRM2 and ODF2 were processed with the VectaFluor™ Duet Immunofluorescence Double Labeling Kit, DyLight® 594 Anti-Rabbit, DyLight® 488 Anti-Mouse (Vector Laboratories; DK-8828) and sections stained with anti-ACTL7B were processed using the Vectastain® ABC-AP Kit (Vector Laboratories; AK-5001) and ImmPACT® Vector® Red alkaline phosphatase substrate (Vector Laboratories; SK-5105). For all stainings an extra 30 min blocking step with 5% BSA in PBS was performed. Fluorescent stainings were DAPI counterstained with ProLong™ Gold antifade reagent (Thermo Fisher Scientific) or ROTI Mount FluorCare DAPI (Carl Roth). Anti-ACTL7B stained sections were counterstained with Haemalum acidic Mayer (Waldeck GmbH & Co KG, Münster, Germany). Stainings were imaged utilizing a Leica DM5500 B microscope (Leica Microsystems, Wetzlar, Germany). Stainings against DYNN1 and DYNN2 were imaged using LSM 710 (Zeiss, Oberkochen, Germany).

### Protein extraction for Immunoblotting and MassSpec analysis

Whole testis tissue was homogenized in 1 ml per 100 mg tissue 1:10 RIPA buffer (Cell Signaling Technology Inc., Danvers, Massachusetts, United States) supplemented with Protease Inhibitor (cOmplete ULTRA Tablets, Mini, EASYpack; Roche, Mannheim, Germany). After incubation on ice for 15 min, the samples were sonicated for 5 min utilizing the Bioruptor UCD-200TM-EX (Tosho Denki Co. LTD., Chiba, Japan). Next, the samples were centrifuged for 30 min at 14,000 rpm at 4°C and the supernatant was used for downstream applications.

### Immunoblotting

Protein extracts were separated on a 12% SDS gel with a 5% stacking gel and transferred to PVDF membranes using the Trans-Blot Turbo System (Bio-Rad). Membranes were blocked using 5% milk for 1 h at room temperature. Primary antibodies (anti-HOOK1 (Proteintech, 10871-1-AP), anti-ACTL7A (Proteintech, 17355-1-AP), anti-ACTL7B (Proteintech, 13537-1-AP), anti-α-tubulin (Santa Cruz Biotechnology, sc-8035), anti-DYNLL1 (Invitrogen, SD08-04), anti-DYNLL2 (Proteintech, 16811-1-AP)) were diluted 1:1000 in 5% milk and membranes were incubated at 4°C overnight. After washing in TBST, the membranes were incubated with secondary antibodies (polyclonal goat anti-rabbit IgG/HRP (P044801-2; 1:2000), polyclonal rabbit anti-mouse IgG/HRP (P026002-2; 1:1000), Agilent Technologies/Dako, Santa Clara, CA, United States) for 1 h at room temperature. Following washing in TBST, the signals were detected using WESTARNOVA2.0 chemiluminescent substrate (Cyanagen) and the ChemiDoc MP Imaging system (Bio-Rad). For western blots after Co-IP SuperSignalTM West Femto Maximum Sensitivity Substrate (Thermo Scientific) was used.

### Macroscopic analysis of testis

Sections of Bouin-fixed testis were processed and stained with Hemalum solution acid (Henricks and Mayer) and Eosin Y solution (Carl Roth, Karlsruhe, Germany) as described (Merges et al., 2022).

### Periodic acid Schiff (Huerta-Cepas et al.) staining

PAS staining was performed as described (Schneider et al., 2020). Deparaffinized, re-hydrated slides were incubated for 10 min in periodic acid (0.5%), washed, incubated 20 min with Schiff reagent and counterstained.

### Isolation of epididymal sperm

Sperm from WT and *Actl7b+/-* males were isolated from the cauda epididymis by swim-out as described (Schneider et al., 2016). Epididymal tissue was incised multiple times and incubated in PBS at 37°C for 20-30 min.

### Transmission electron microscopy

Testis tissue was prepared as described (Merges et al., 2022). In brief, the tissue was fixed in 3% glutaraldehyde at 4°C overnight, washed, post-fixed with 2 % osmium tetroxide at 4°C for 2 h and again washed. After dehydration and contrasting in 70% (v/v) ethanol 0.5% (m/v) uranyl acetate (1 – 1.5 h, 4°C), samples were washed with propylenoxide (3x 10 min, RT) and stored in propylenoxide:Epon C (1:1, (v/v)) at 4°C overnight. Next, the pellets were embedded in Epon C (70 °C, 48 h) and ultra-thin sections were prepared. Ultrathin sections were contrasted with UranyLess (Electron Microscopy Sciences, Hatfield, PA, USA) and lead citrate. Images were taken using the Philips CM10 transmission electron microscope equipped with analySiS imaging software and Zeiss Crossbeam 550 FIB SEM equipped with a retractable STEM detector.

### Eosin-Nigrosin staining

Staining was performed as described (Merges et al., 2022) using 50 μl of sperm swim-out and 50 μl Eosin-Nigrosin stain (0.67 g Eosin Y (color index 45380), 0.9 g sodium chloride, 10 g Nigrosin (color index 50420), 100 ml ddH_2_O). 200 sperm per animal were analyzed.

### Co-immunoprecipitation (Co-IP)

Proteins from WT C57BL/6J testis were isolated utilizing the T-PER™ Tissue Protein Extraction Reagent (Thermo Fisher Scientific) supplemented with Halt Protease Inhibitor Single-Use Cocktail EDTA-Free (Thermo Fisher Scientific) according to the manufacturers instructions.

The ACTL7B antibody (Proteintech; 13537-1-AP) was purified using the Amicon® Ultra 30K – 0.5 Centrifugal Filter Device (Merck Millipore Ltd., Burlington, MA, USA; UFC503008) according to the manufacturer’s instructions. The purified antibody was coupled to beads using the Dynabeads™ Antibody Coupling Kit (Thermo Fisher Scientific; 14311D), using 7µg antibody per mg beads. Next, 7.5 mg of antibody-coupled beads and 5 mg empty beads were used with the Dynabeads™ Co-Immunoprecipitation Kit (Thermo Fisher Scientific; 14321D) according to the manufacturer’s instructions. Proteins eluted from the beads in 1 ml HPH EB (0.5 M NH_4_OH, 0.5 mM EDTA) buffer. 700 µl were lyophilized and sent for mass spec analysis. 300 µl were lyophilized and solubilized in SDS-PAGE sample loading buffer.

DYNLL1 and DYNLL2 antibodies were also purified using Amicon® Ultra 30K – 0.5 Centrifugal Filter Device (Merck Millipore Ltd). For antibody coupling using the Dynabeads™ Antibody Coupling Kit (Thermo Fisher Scientific) 10 µg antibody per mg beads were used. 1.5 mg beads were used with the Dynabeads™ Co-Immunoprecipitation Kit (Thermo Fisher Scientific) for WesternBlot applications according to the manufacturer’s instructions.

### MassSpec

Proteins from whole testis from five WT, Actl7b+/– and Actl7b–/– mice were isolated as described and used for mass spectrometric analysis. Peptide preparation and liquid chromatography (LC)-mass spectrometry (MS) were performed at the University of Bonn Core Facility Mass Spectrometry.

### Preparation of co-immunoprecipitation samples for LC/MS

50 µg of protein per sample were subjected to in solution preparation of peptides with the iST 96x sample preparation kit (Preomics GmbH, Martinsried, Germany) according to manufacturer’s recommendations.

### Whole cell proteomic analysis: precipitation, proteolysis, peptide labeling, and fractionation

Protein lysates with 200 µg protein were mixed with a 4-fold volume of chilled acetone (−20°C). After 1 h at -20°C proteins were sedimented by centrifugation for 15 min at 14,000xg. The supernatant was discarded und pellets air-dried. Pellets were dissolved in 20 µL Lyse buffer (Preomics iST-NHS kit) and protein content determined by BCA assay. Solutions with 40 µg protein were mixed with 50 µL of the kit’s DIGEST solution (3h, 37°C). 0.25 mg of TMTpro isobaric Mass Tag Labeling reagent (15plex) were added to each sample and incubated at room temperature for 1 h. 10 µL 5% hydroxylamine were used to quench the reaction. The preparation procedure was continued according to the iST-NHS kit instructions. Pooled peptides were dried in a vacuum concentrator, dissolved in 20 mM ammonium formate (pH=10) and fractionated by reversed phase chromatography at elevated pH with a Reprosil 100 C18 column (3 µm 125 × 4 mm, Dr. Maisch GmbH, Ammerbuch-Entringen, Germany). 60 fractions were combined into 6 pools and dried in a vacuum concentrator. Peptides were purified by solid phase extraction (Oasis HLB cartridges, Waters GmbH, Eschborn, Germany).

### LC/MS measurements

Before measurement peptides were re-dissolved in 0.1% formic acid (FA) to yield a 1 g/L solution and separated on a Dionex Ultimate 3000 RSLC nano HPLC system (Dionex GmbH, Idstein, Germany). The autosampler was operated in μL-pickup mode. 1 µL was injected onto a C18 analytical column (self-packed 400 mm length, 75 µm inner diameter, ReproSil-Pur 120 C18-AQ, 1.9 µm, Dr. Maisch). Peptides were separated during a linear gradient from 5% to 35% solvent B (90% acetonitrile, 0.1% FA) at 300 nL/min. The nanoHPLC was coupled online to an Orbitrap Fusion Lumos mass spectrometer (ThermoFisher Scientific, Bremen, Germany).

#### Measurement of peptides from co-immunoprecipitated proteins

Gradient length was 90 min. Peptide ions between 300 and 1600 m/z were scanned in the Orbitrap detector every 3 s with a resolution of 120,000 (maximum fill time 50 ms, AGC target 100%). Polysiloxane (445.12002 Da) was used for internal calibration (typical mass error ≤1.5 ppm). In a top-speed method, peptides were subjected to higher energy collision induced dissociation (1.0 Da isolation, normalized energy 27%) and fragments analyzed in the Orbitrap (resolution 15,000, AGC target 100%, maximum fill time 22 ms). Fragmented peptide ions were excluded from repeat analysis for 20 s.

#### Measurement of TMT-labelled fractions

Gradient length was 150 min. Peptide ions between 330 and 1600 m/z were scanned in the Orbitrap detector with settings as above. In a top-speed method, peptides were subjected to collision induced dissociation for identification (CID: 0.7 Da isolation, normalized energy 30%) and fragments analyzed in the linear ion trap with AGC target 50% and maximum fill time 35 ms, rapid mode. Fragmented peptide ions were excluded from repeat analysis for 30 s. Top 10 fragment ions were chosen for synchronous precursor selection and fragmented with higher energy CID (HCD: 3 Da MS2 isolation, 65% collision energy) for detection of reporter ions in the Orbitrap analyzer (range 100-180 m/z, resolution 50,000, maximum fill time 86 ms, AGC target 200%).

### Data analysis

Raw data processing and database search were performed with Proteome Discoverer software 2.5.0.400 (Thermo Fisher Scientific). Peptide identification was done with an in-house Mascot server version 2.8.1 (Matrix Science Ltd, London, UK). MS data were searched against Mus musculus sequences from the SwissProt database including isoforms (2022/03, 17132 murine sequences) and contaminants database (cRA (Mellacheruvu et al., 2013)). Precursor Ion m/z tolerance was 10 ppm, fragment ion tolerance 0.5 Da (CID). Tryptic peptides with up to two missed cleavages were searched. C_6_H_11_NO-modification of cysteines (delta mass of 113.08406) and TMTpro on N-termini and lysines were set as static modifications. Oxidation was allowed as dynamic modification of methionine. Mascot results were evaluated by the Percolator algorithm version 3.02.1(The et al., 2016) as implemented in Proteome Discoverer. Spectra with identifications above 1% q-value were sent to a second round of database search with semi tryptic enzyme specificity (one missed cleavage allowed). Protein N-terminal acetylation, methionine oxidation, TMTpro, and cysteine alkylation were then set as dynamic modifications. Actual FDR values were 0.7% (peptide spectrum matches) and 1.0% (peptides and proteins). Reporter ion intensities (most confident centroid) were extracted from the MS3 level, with SPS mass match >65%.

### Differential abundance analysis

Data for proteins detected in all genotypes and replicates were with more than 2 peptides were log2 transformed and median normalized. Abundances were analyzed as described before (Merges et al., 2022). The Bioconductor package proDA (Ahlmann-Eltze and Anders, 2021) was used (peptide spectrum match (PSM)-level data extracted from Protein Discoverer). Proteins with LFC>0.5 and LCF>1 (FDR adjusted p<0.05) compared to WT were analyzed. The R-package ggplot2 (Wickham, 2011) was used to plot the data.

### F-/G-Actin ratio

The F-/G-Actin ratio was analyzed as described (Pilo Boyl et al., 2007). Whole testis tissue was homogenized in 500 µm 1.1 x PHEM buffer (600 mM PIPES-Na, 250 mM HEPES-Na, 100 mM EGTA, 20 mM MgCl_2_, pH 6.9) with 1.2 % TritonX-100 and incubated on ice for 15 min. Next, the samples were centrifuged in a swing-out rotor for 10 min at 10000 g at 4°C. 400 µl of the G-actin congaing supernatant were boiled with 100 µl of 5x SDS loading buffer (110 mM Tris/HCl pH 6.8, 20% Glycerol, 3.8% SDS, 8% β-mercaptoethanol, 0.05% bromophenol blue) for 10 min and then cooled on ice. After discarding the rest of supernatant, the pellet was dried and resuspended in 650 µl 1x SDS loading buffer, boiled for 10 min and cooled on ice. Equal volumes of supernatant and pellet fraction were loaded on a 10% acrylamide gel, the gel was blotted on methanol-activated PVDF membrane and anti-β-actin (1:5000; 08-691002; MP Biomedicals, Santa Ana, CA, USA) was used to detect the G- and the F-actin fractions. Band quantification was performed using the GE FujiFilm ImageQuant LAS-4000 CH mini Imager (GE Healthcare, Chicago, IL, USA).

### Evolutionary analysis

Evolutionary rates of *ACTL7B* among rodents and primates were analyzed according to Lüke et al. (Lüke et al., 2016). *ACTL7B* coding sequences were obtained from NCBI genbank and Ensembl genome browser. Phylogenetic trees of considered species were built according to the “Tree of Life web project” (Letunic and Bork, 2021). The webPRANK software was applied for codon-based phylogeny-aware alignment of orthologous gene sequences (Loytynoja and Goldman, 2010). The tree and alignment were visualized using the ETE toolkit (Huerta-Cepas et al., 2016).

To evolutionary rates and selective pressures were determined using codeML implemented in PAML4.9 (Yang, 1997, 2007). The evolutionary rate is based on the calculation of the nonsynonymous/synonymous substitution rate ratio (ω = dN/dS). It distinguishes between purifying selection (ω<1), neutral evolution (ω=1) and positive selection (ω>1).

Different null and alternative models (M) were applied. The M0 model served as basic model for all performed analyses. Different codon frequency settings were tested for the M0 model of each gene and the setting with the highest likelihood was chosen. To test whether alternative models describe the selective constraints within a dataset better than the null models, likelihood-ratio-tests (LRT) were performed.

In order to determine the overall evolutionary rate and selective pressure on the coding sequence among all included species we employed two models: M0 “one ratio” in which all branches were constrained to evolve at the same freely estimated evolutionary rate; M0fix (fixed ratio) in which the evolutionary rate for all branches was constrained to 1. The M0 model calculates the overall evolutionary rate. An LRT between M0 and M0fix was performed to determine of the calculated evolutionary rate significantly differs from 1 (neutral).

In order to determine if the selective pressures differ between rodents and primates we employed two models: M0 “one ratio”; MCfree “two-ratio” which allows the estimation of a free and independent ω for the two marked clades. To test if the alternative MCfree model presents a better fit for the data we compared the models log likelihood values by LRT.

To test evolution along coding sequences and infer codon sites under positive or purifying selection we applied LRT comparing the null model M1a “nearly neutral”, which does not allow sites with ω>1 with the alternative model M2a “selection” which does. The models assigns the codon sites into different classes: Class 0: sites under purifying selection (0>ω>1); Class 1: sites evolving neutrally (ω = 1); Class 2a (only M2a): sites subject to positive selection (ω<1). Bayesian statistics were used to identify those codons that have been subject to either positive selection (if the alternative model is the better fit) or purifying selection (if the null model is the better fit). Only sites with posterior probabilities (Bayes Empirical Bayes) higher than 0.95 to be assigned to class 0 or class 2a were determined to be under purifying or positive selection respectively.

### Statistics

Values are, if not indicated otherwise, given as mean values with standard deviation. Statistical significance was calculated using the two-tailed, unpaired Student’s t-test. Values of p < 0.05 were considered significant (p < 0.05= *; p < 0.005= **; p < 0.001= ***).

## Competing interests

The authors declare no competing or financial interests.

## Acknowledgements

We thank Gaby Beine, Greta Zech, Andrea Jäger, Angela Egert and Irina Kosterin for excellent technical assistance. Protein identification was performed at the University of Bonn Core Facility Mass Spectrometry, Institute of Biochemistry and Molecular Biology, Medical Faculty, University of Bonn funded by the Deutsche Forschungsgemeinschaft (DFG, German Research Foundation) – Projektnummer 386936527. We thank the Microscopy Core Facility of the Medical Faculty at the University of Bonn for providing support and instrumentation funded by the DFG (388171357).

## Funding

This study was supported by grants from the Deutsche Forschungsgemeinschaft (DFG) to HS (SCHO 503/27-1)

## Author contributions

Conceptualization: H.S.; Methodology: G.E.M., L.A.; Formal analysis: G.E.M., L.A., D.G.d.R., M.J., W.W.; Investigation: G.E.M., L.A., K.L.; Resources: H.S.; Data curation: L.A.; Writing - original draft: G.E.M.; Writing - review & editing: G.E.M., L.A., H.S.; Visualization: G.E.M., L.A.; Supervision: H.S.; Project administration: H.S.; Funding acquisition: H.S.

## Notes

### Competing Interest Statement

The authors have declared no competing interest.

