## Supplementary figures and images for "*Actl7b*-deficiency leads to mislocalization of LC8 type dynein light chains and disruption of murine spermatogenesis"

### Figure S1

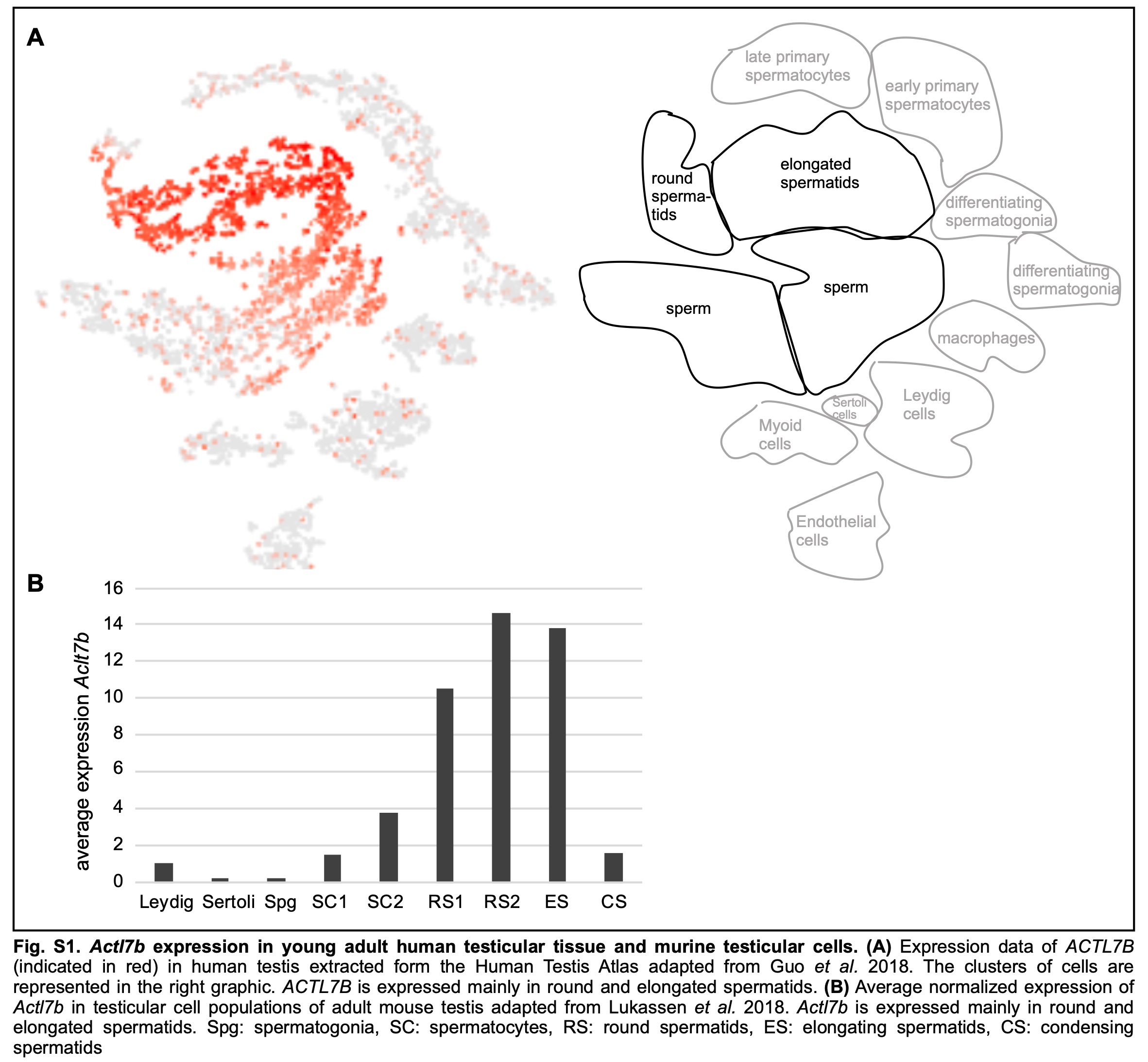

### Figure S2

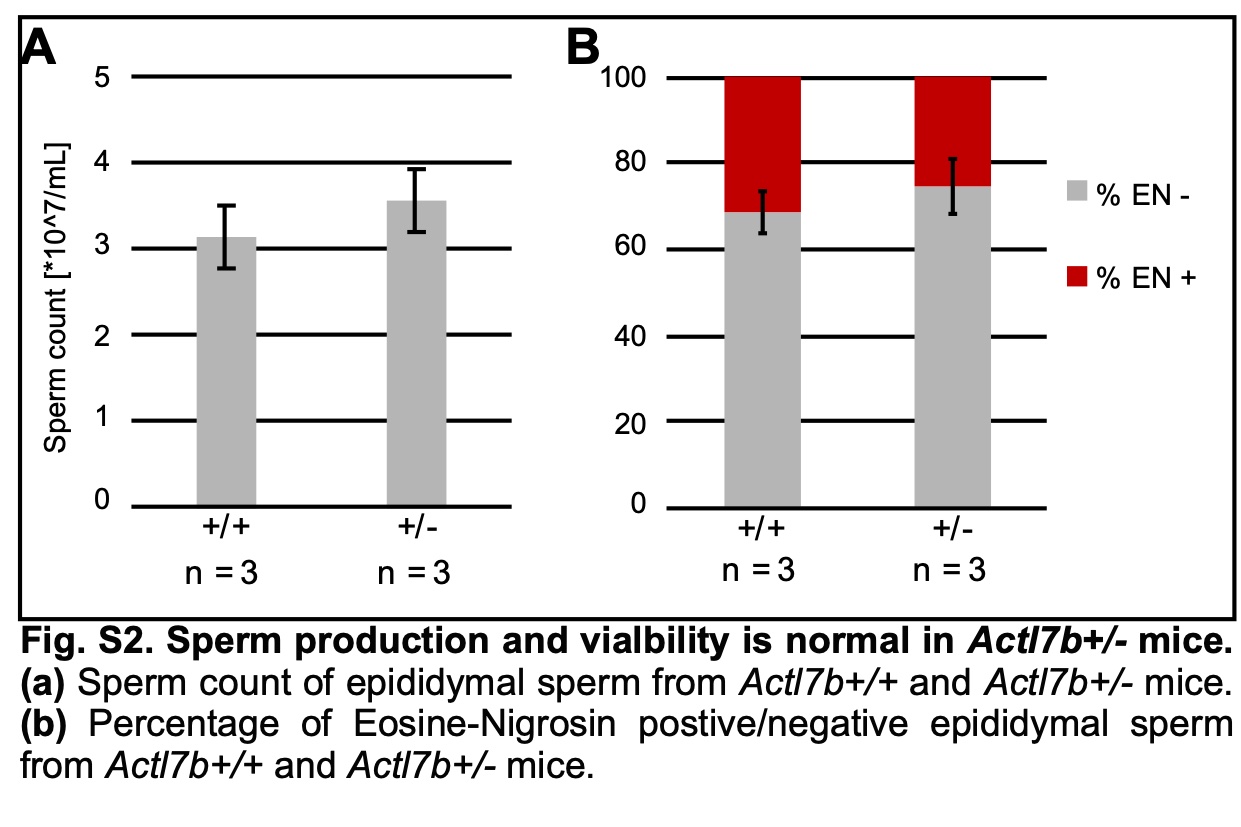

### Figure S3

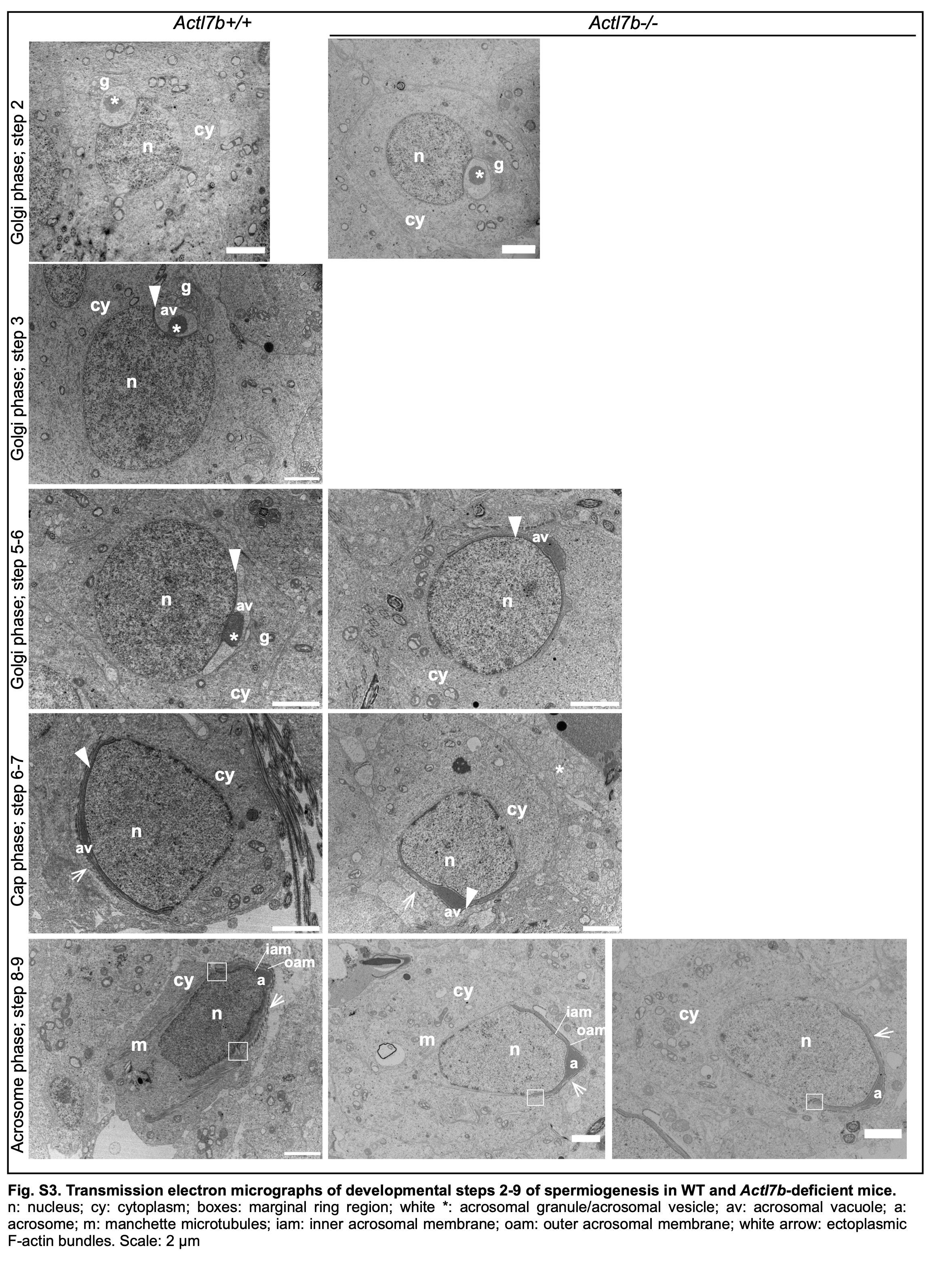

### Figure S4

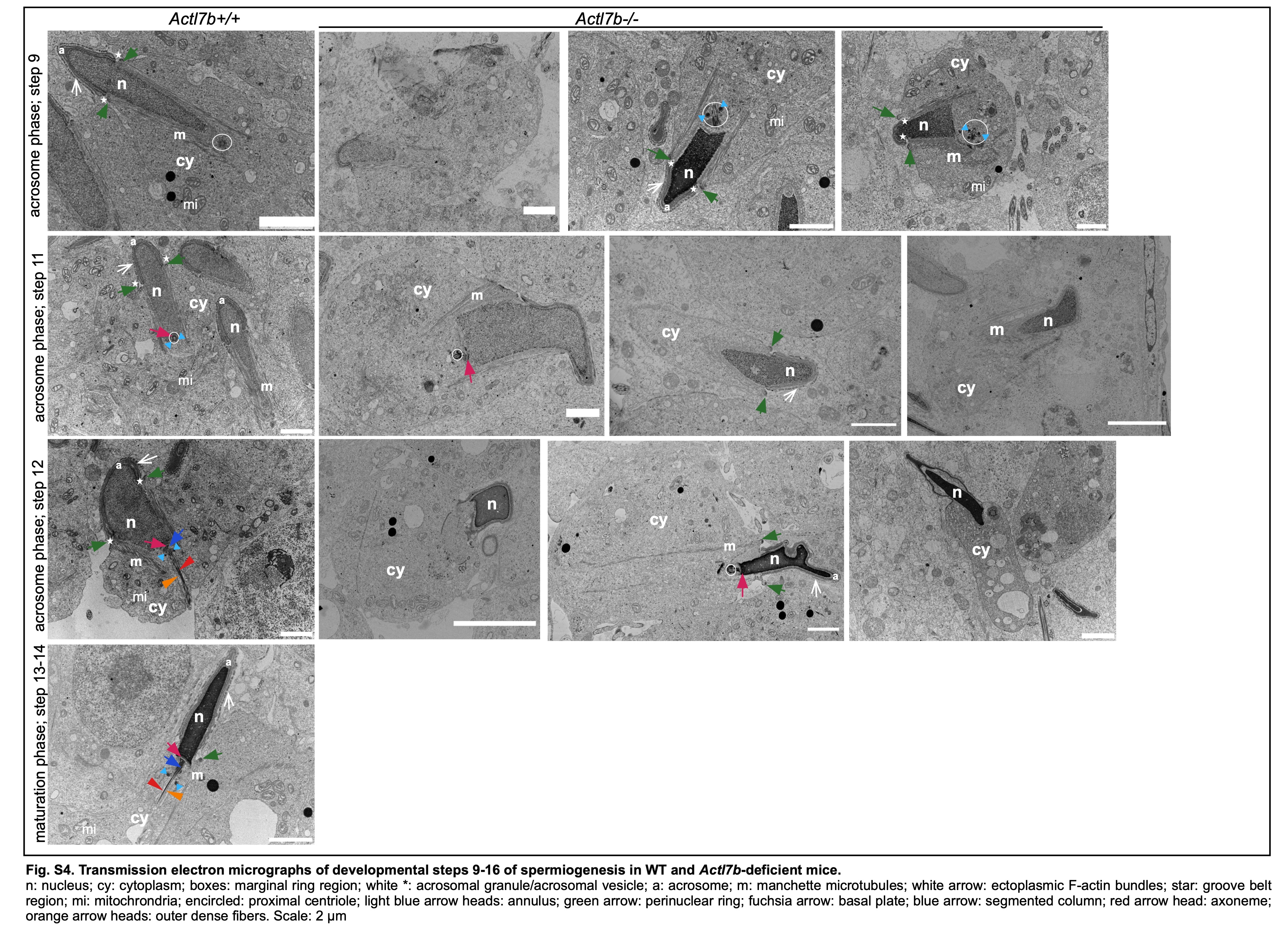

### Figure S5

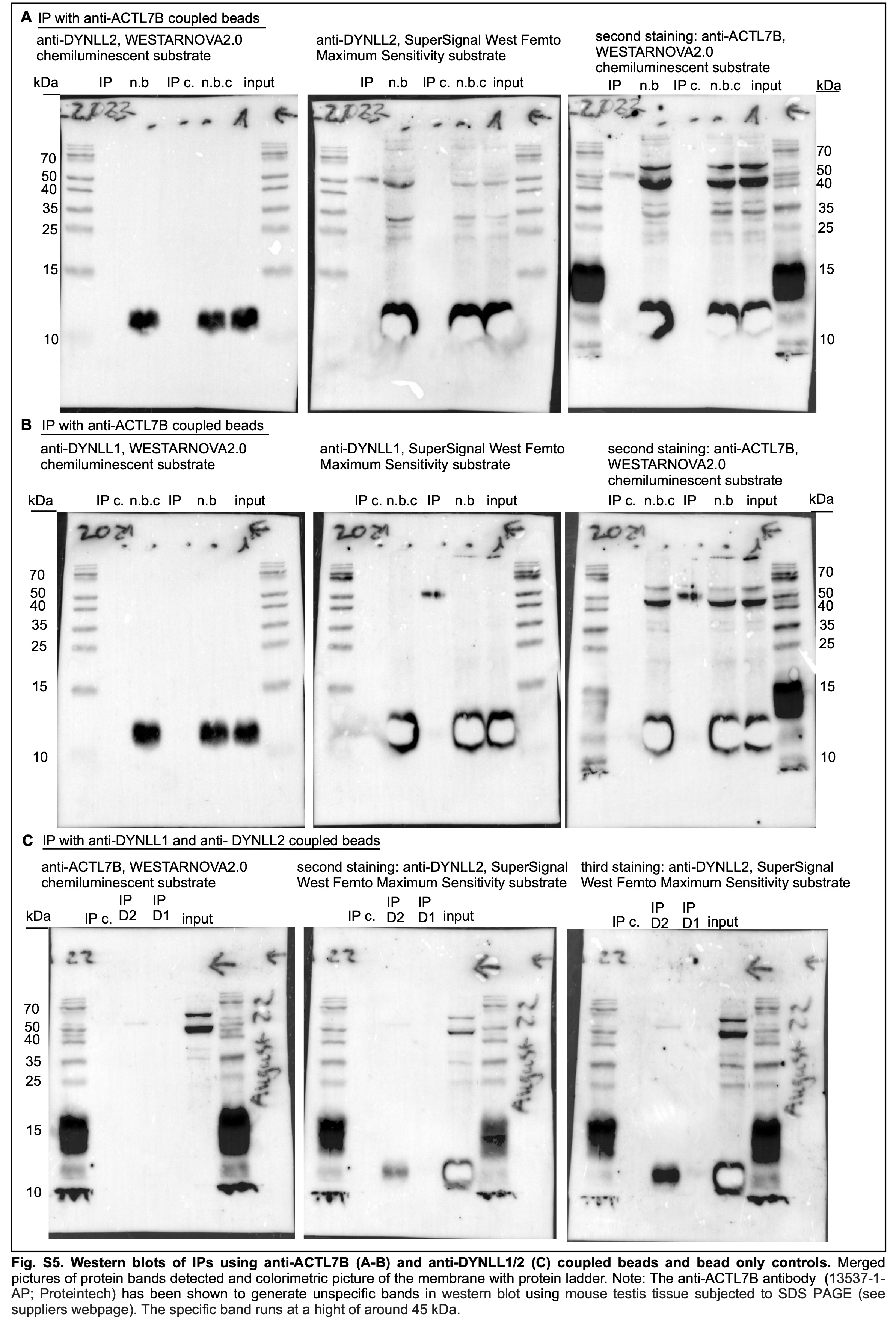

### Figure S6

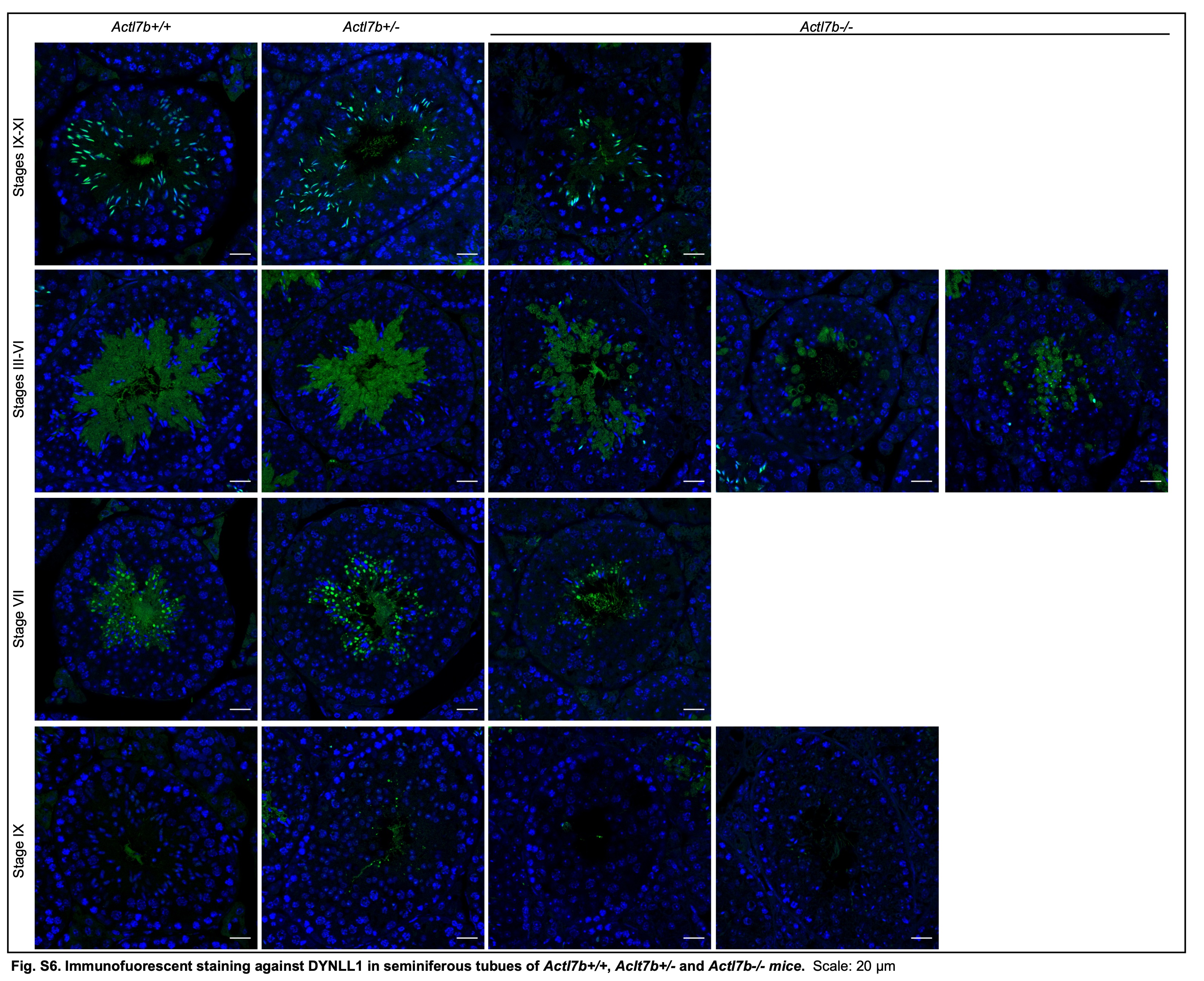

### Figure S7

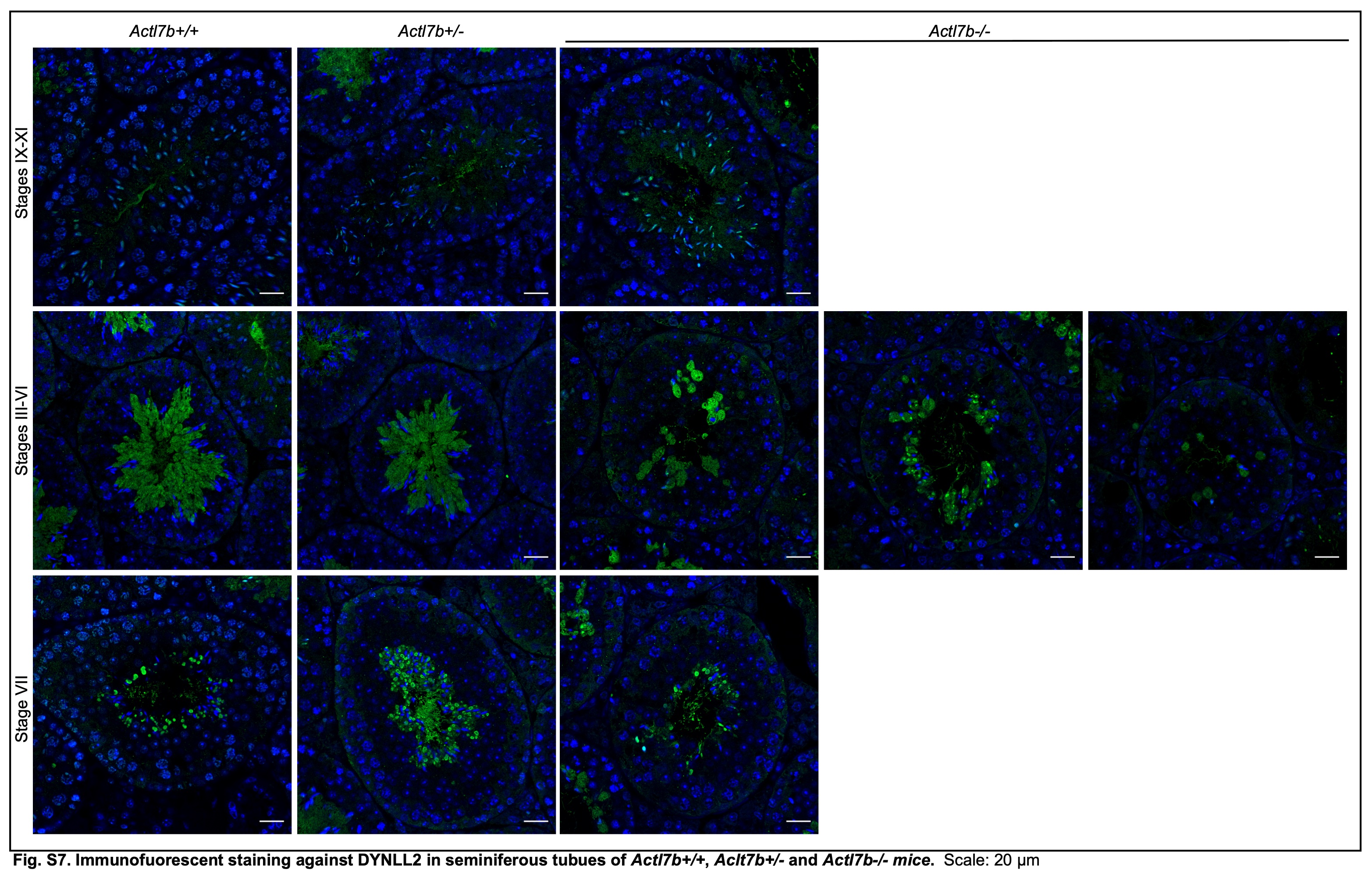

### Figure S8

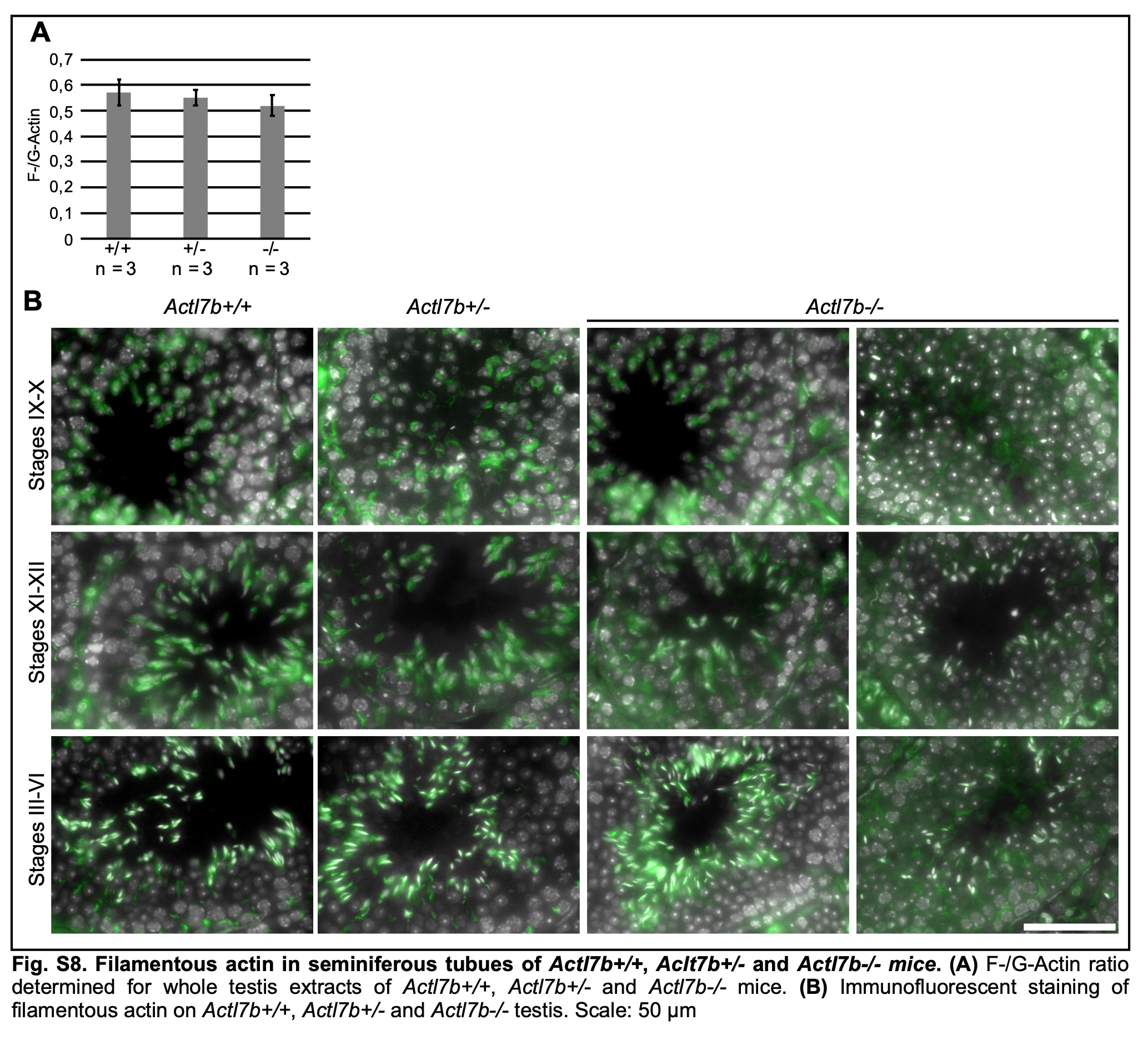

### Figure S9

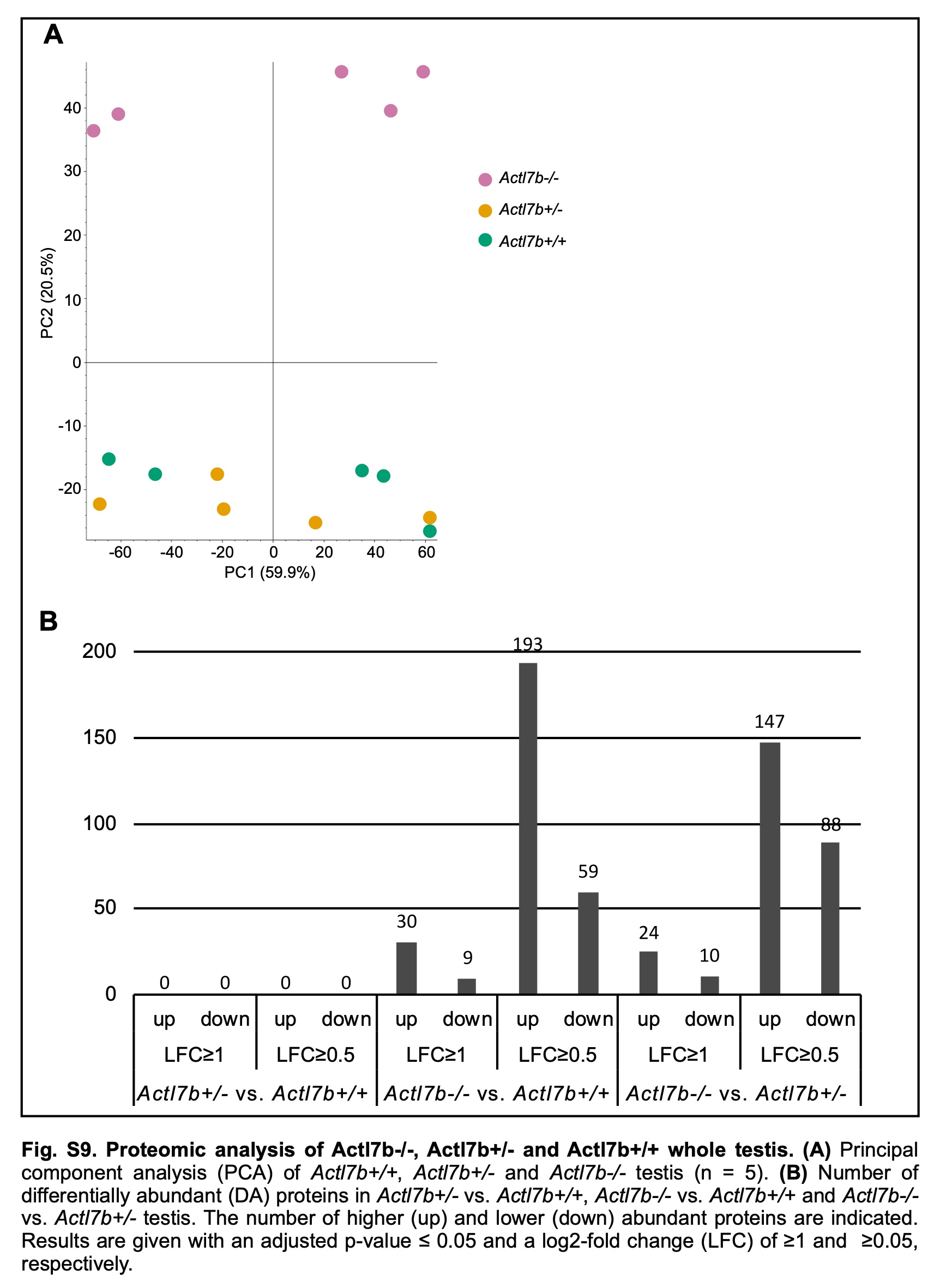

### Figure S10

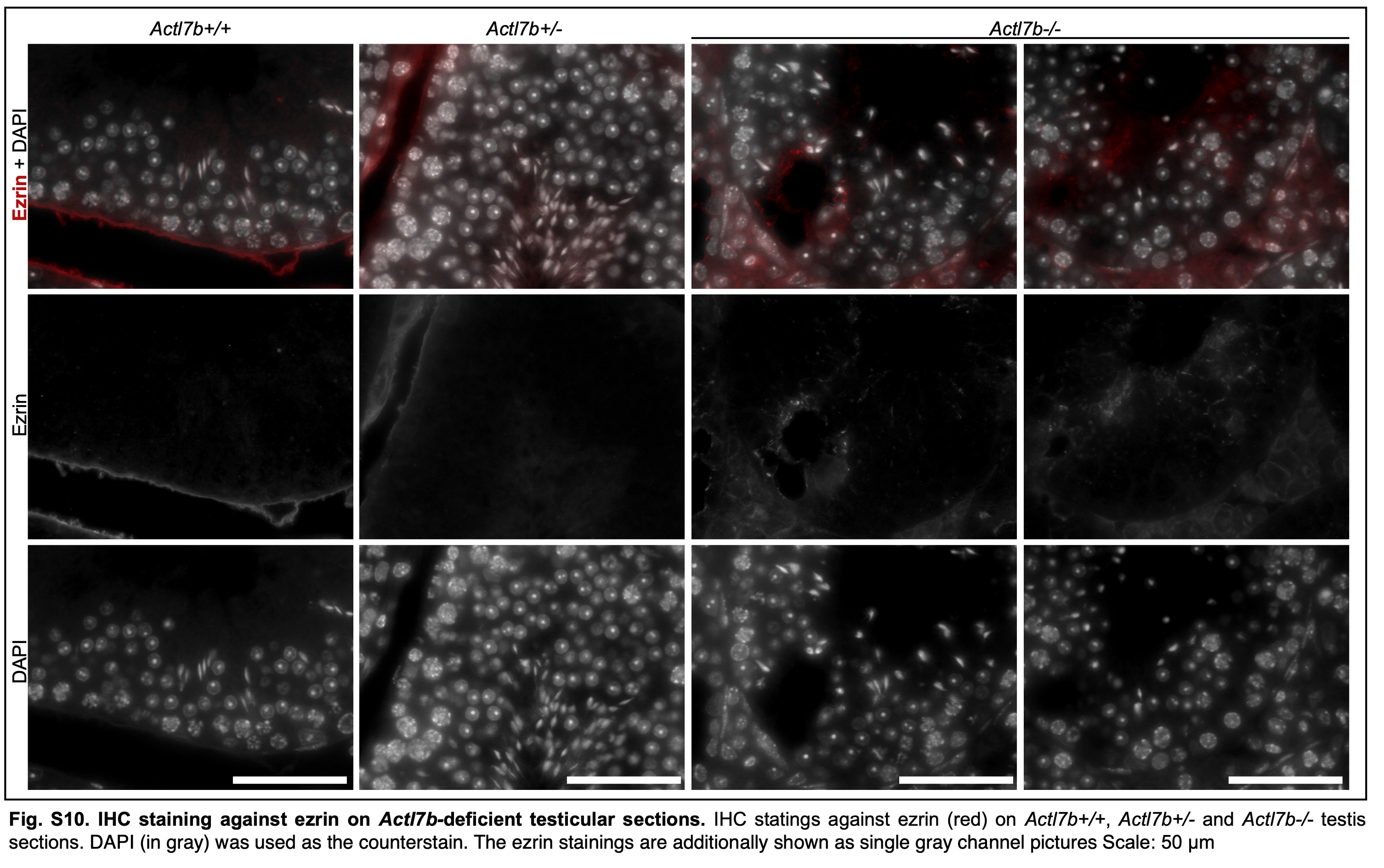

### Figure S11

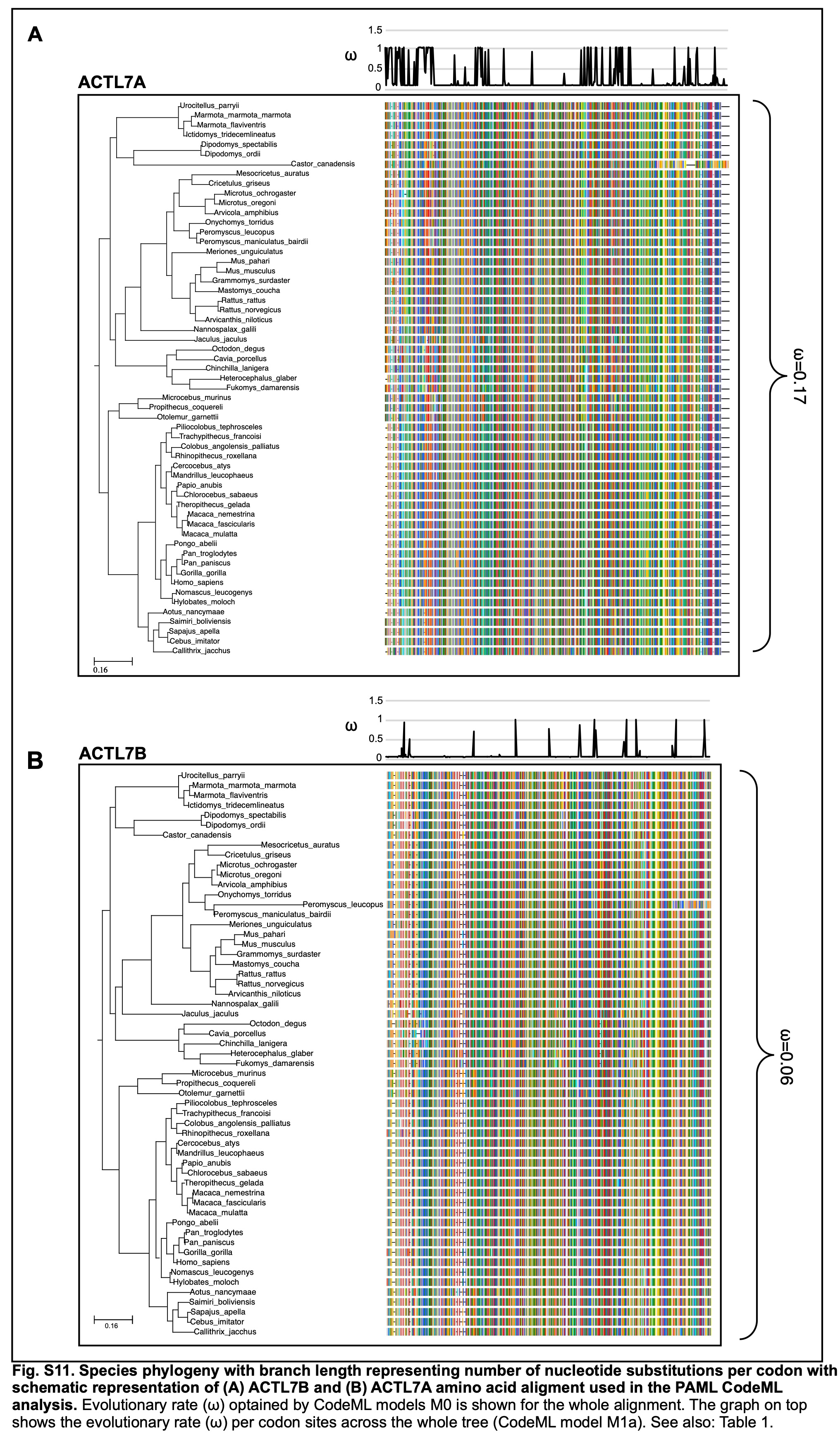

### Table S1

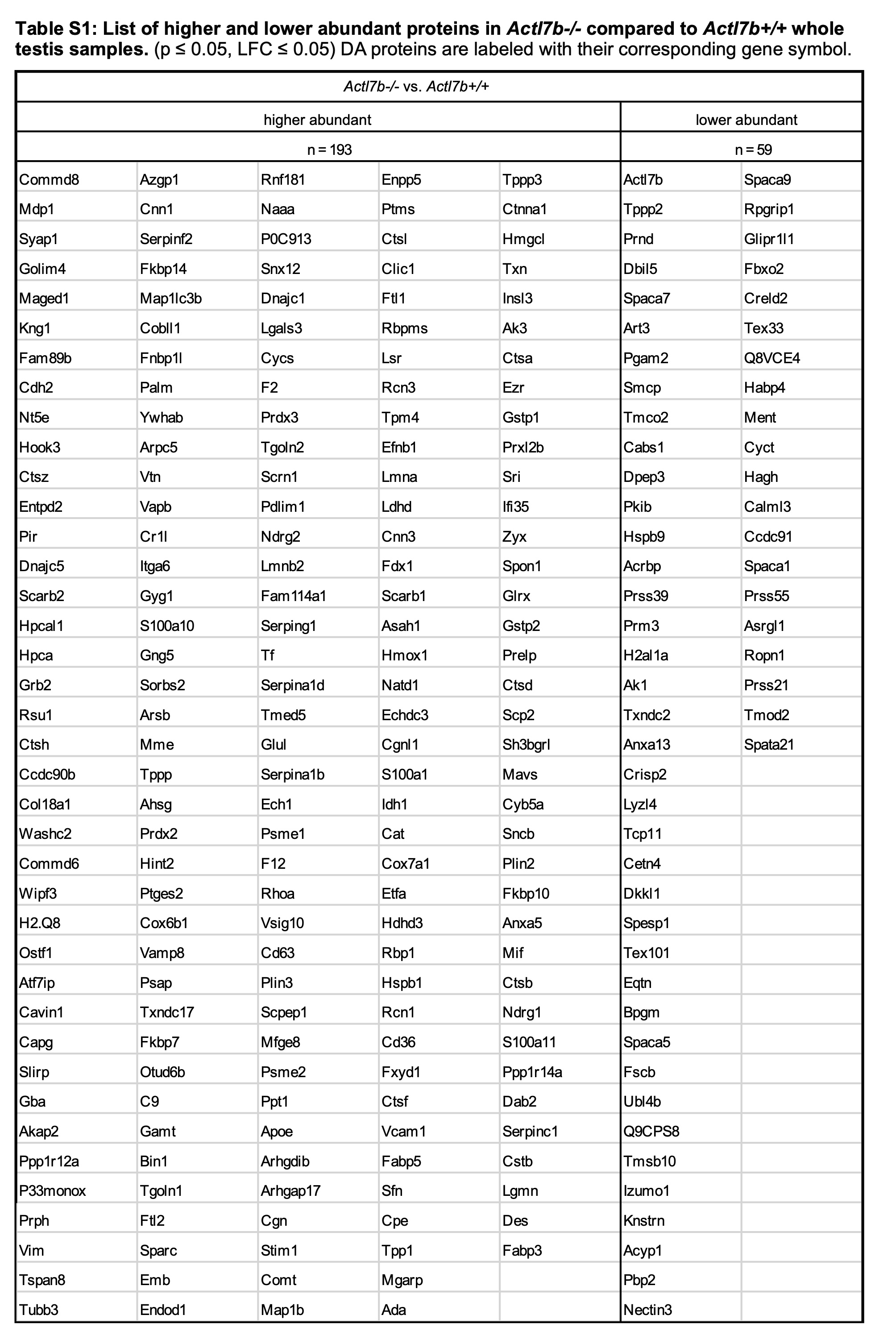
